# Imaging FCS Delineates Subtle Heterogeneity in Plasma Membranes of Resting Mast Cells

**DOI:** 10.1101/794248

**Authors:** Nirmalya Bag, David A. Holowka, Barbara A. Baird

## Abstract

A myriad of transient, nanoscopic lipid- and protein-based interactions confer a steady-state organization of plasma membrane in resting cells that is poised to orchestrate assembly of key signaling components upon reception of an extracellular stimulus. Although difficult to observe directly in live cells, these subtle interactions can be discerned by their impact on the diffusion of membrane constituents. Herein, we quantified the diffusion properties of a panel of structurally distinct lipid-anchored and transmembrane (TM) probes in RBL mast cells by multiplexed Imaging Fluorescence Correlation Spectroscopy. We developed a statistical analysis of data combined from many pixels over multiple cells to characterize differences as small as 10% in diffusion coefficients, which reflect differences in underlying interactions. We found that the distinctive diffusion properties of lipid-anchored probes can be explained by their dynamic partitioning into ordered proteo-lipid nanodomains, which encompass a major fraction of the membrane and whose physical properties are influenced by actin polymerization. Effects on diffusion by functional protein modules in both lipid-anchored and TM probes reflect additional complexity in steady-state membrane organization. The contrast we observe between different probes diffusing through the same membrane milieu represent the dynamic resting steady-state, which serves as a baseline for monitoring plasma membrane remodeling that occurs upon stimulation.

## INTRODUCTION

Cells typically exist in noisy environments, and their plasma membranes are poised to respond optimally to external chemical and physical stimuli, including specific chemical ligands (1), thermal shock (2), and electrical and mechanical forces (3, 4). For versatile and efficient responses, the membrane accommodates receptors and other structures that sense the external stimuli as well as organizes surrounding lipids and proteins (5). Many of the underlying interactions are cooperative and weak, providing a dynamic steady-state platform that has the capacity to respond to a specific stimulus over environmental noise and to regulate transmembrane signaling components (6, 7). Key to responsive membrane organization are structural configurations that can be modulated to selectively include/exclude other components. A prominent example is “lipid rafts,” an ill-defined term for dynamic proteolipid nanodomains that resemble the liquid ordered (Lo) phase in model membranes (8). Although there is ample experimental support for participation of these Lo-like proteolipid nanodomains (a term we currently prefer to “rafts”) in stimulated signaling (7, 9–13), their physical nature has been difficult to define because of their diversity, their sub-resolution dimensions, and their transience (12–19). Over the years we have used a wide range of approaches to examine these nanodomains because of their clear participation in transmembrane signaling initiated by the high affinity receptor (FcεRI) for immunoglobulin E (IgE) on mast cells, as triggered by multivalent antigen in the allergic immune response (20). Cells are sensitized to antigen when IgE antibodies bind to FcεRI, which diffuse as monomeric species in the plasma membrane (21, 22). Cell activation occurs only after addition of antigen, which crosslinks the IgE-FcεRI to stabilize their association with Lo-like proteolipid nanodomains and consequently their functional coupling with Lyn tyrosine kinase. The stimulatory event depends on shifting the balance of phosphorylation-dephosphorylation of cross-linked IgE-FcεRI towards phosphorylation, leading to downstream signaling. This is facilitated by the capacity of the nanodomains to preferentially include the key kinase (Lyn), which is anchored to the inner leaflet by Lo-preferring fatty acid chains, and exclude a transmembrane phosphatase, which is accommodated more favorably in a liquid disordered, Ld-like environment (23). This example illustrates the predisposition of the plasma membrane to respond to a specific stimulus by the steady state presence of Lo-like proteolipid nanodomains that are dynamic in nature but are stabilized and utilized when the stimulus arrives.

Co-existence of membrane structures that facilitate spatial compartmentalization underlies the hierarchal model proposed by Kusumi and colleagues, based primarily on their extensive ultrahigh-speed single particle tracking (SPT) and scanning electron microscopy measurements, with additional features drawn from complementary studies in other laboratories (Figure 1A) (24–29). The hierarchal model builds on membrane compartments (corrals; 40–230 nm) defined by the long chain actin meshwork (fence) with anchored transmembrane proteins (pickets). Importantly, such actin-based compartmentalization imposes fundamental membrane organization by preventing Lo/Ld lipid phase separation (30–33). Considerable evidence supports the view that nanoscale Lo-like channels align along the picket fences, either by the effects of critical behavior and pinned Lo-preferring components (30, 31, 34) or by stabilization with Lo-preferring protein pickets (35, 36). In the hierarchical model, Lo-like nanodomains (2–20 nm) and protein complexes (3–10 nm) also exist within the corrals. In addition, experiments and simulations underlying the active composite model of Mayor, Rao, and colleagues showed that ordered lipid nanodomains also arise from active, myosin-driven asters of short actin chains that connect to inner leaflet lipids and cause alignment of tails of Lo-preferring lipids in the outer leaflet (37–39).

**Figure 1.**
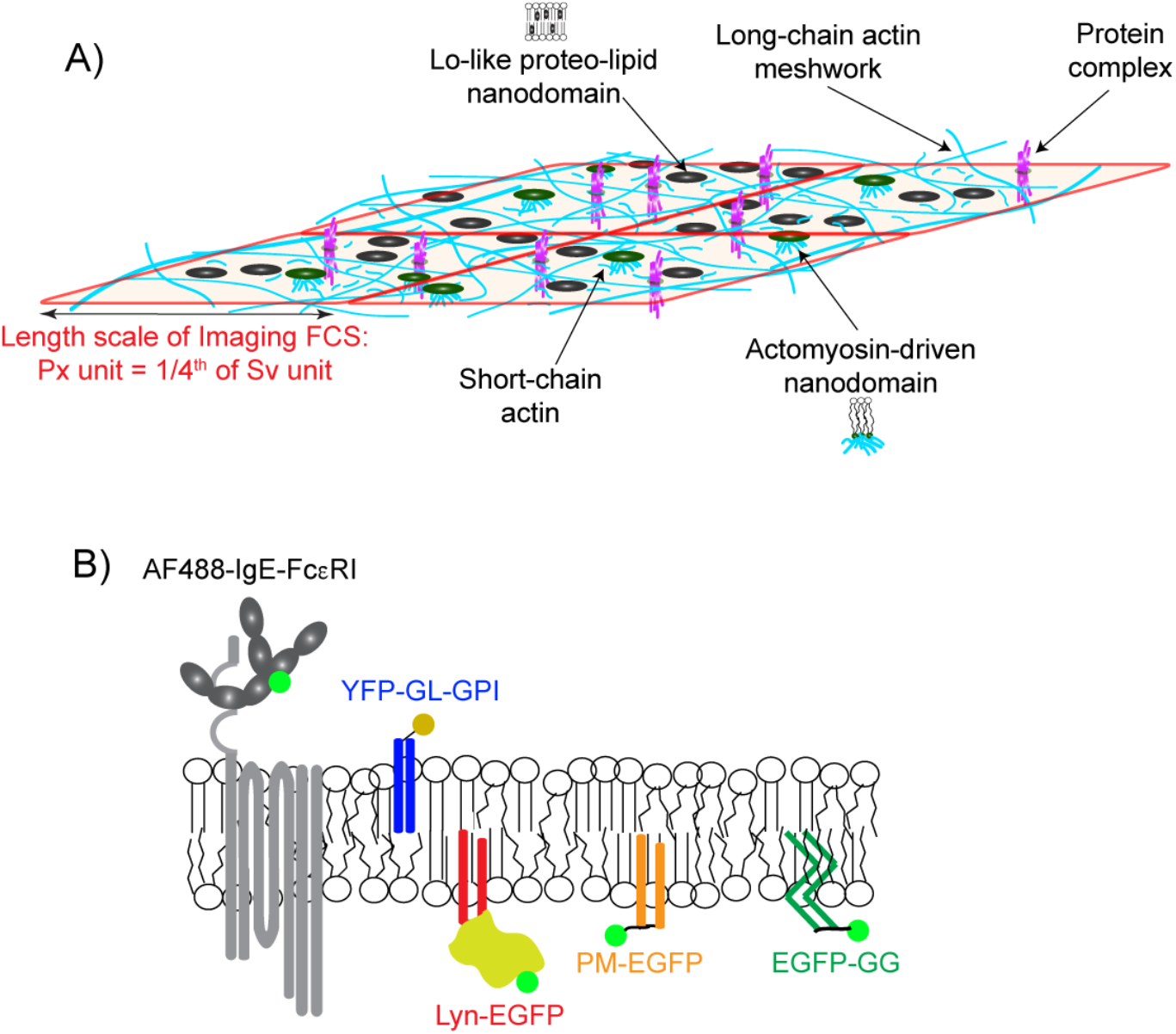
The composite plasma membrane organization is evaluated by monitoring the diffusion of structurally distinct probes. **A**) The plasma membrane is organized at different length scales in a hierarchical scheme: a relatively static actin meshwork (cyan long chains); dynamic Lo-like proteolipid nanodomains (black circles) with variable physical properties within and across leaflets; transmembrane ordered lipids mediated by dynamic, myosin-driven assembly of short actin chains (green circles connected to short cyan chains); stable or dynamic protein complexes. We note that lipid nanodomains and protein complexes are much smaller than the dimensions of the actin meshwork and not drawn to scale here. Interactions of a probe with these organizational features retards its diffusion, depending on that probe’s physico-chemical properties. ImFCS measures diffusion coefficients (*D*) at the resolution of a Px unit which has dimensions considerably larger than the actin meshwork, as described in text; parameters *τ*_*0*_ and 1/*D*_*eff*_ = *Slope* are measured in Sv units which comprise a square of 16 Px units. **B**) Fluorescently-labeled, lipid-anchored and transmembrane probes evaluated in this study.

Direct imaging of dynamic plasma membrane heterogeneity at the nanoscale is challenging, even with super-resolution optical microcopy (40, 41). Fluorescence spectroscopies, which are often coupled with diffraction-limited microcopy, offer new possibilities to extract dynamic properties of plasma membranes (42). As described herein, we employ Imaging Fluorescence Correlation Spectroscopy (ImFCS) (43), a camera-based modality of FCS (44), to examine the effects of membrane organization on diffusion of individual membrane components, based on the premise that local structures, mediated by proteins or lipids or both, curtails lateral diffusion differentially (41, 45, 46). Within the extended hierarchical model described above, we take the basic view that the plasma membrane comprises Lo-like proteolipid nanodomains (along and within the boundaries of corrals) connected by Ld-like regions, and that this organization is modulated by long and short chain actin (Figure 1A). With ImFCS we can quantify diffusion of multiple, diverse probes that differentially interact with membrane constituents and integrate these distinctive diffusion behaviors to create a composite picture of the plasma membrane organization.

Compared to conventional single-spot FCS measurements, the family of image-based fluctuation methods offers various kinds of spatial analysis in addition to evaluation of diffusion. The spatio-temporal information obtained from these methods depends on the resolution of the microscope, scanning configuration and modes of fluctuation analysis (47–49). The ImFCS we employ was developed as an ensemble-averaged but single molecule sensitive technique, which provides a pixelated map of membrane diffusion (43, 50). A continuous series of total internal reflection fluorescence microscopy (TIRFM) images of the ventral plane of fluorescently labeled, live cells is captured by a fast, sensitive camera, which spatially divides the image into an array of submicron pixels (49, 50). Each pixel of the camera corresponds to a diffraction-limited membrane spot. Autocorrelation function (ACF) analysis of temporal fluorescence fluctuations of each pixel yields a macroscopic, Brownian diffusion coefficient (*D*) at that pixel. ImFCS data acquisition for a single cell typically contains hundreds of pixels, such that hundreds of parallel FCS experiments are carried out on that cell. When the pixel measurements for a single cell, or for multiple cells, can be combined into an ensemble, ImFCS can deliver much more robust estimates of diffusion compared to conventional FCS that is based on a single illumination volume. ImFCS addresses limitations in spatial resolution using spot variation FCS (svFCS), as developed by Lenne and colleagues for conventional FCS, which indirectly detects the existence of sub-resolution regions of confined diffusion (51, 52). In an ImFCS experiment, the fluorescence fluctuations collected for each pixel can be used directly to perform svFCS analysis because pixel binning (i.e, summing over adjacent pixels) effectively generates spot areas of variable sizes (53). Incorporating inherent multiplexing capacity, straightforward implementation, and compatibility with a conventional live cell imaging platform, ImFCS offers enhanced capabilities for directly evaluating macroscopic diffusion properties and indirectly assessing possible influence of sub-resolution domains of confinement (50).

Our goal in this study is to gain a detailed knowledge of spatio-temporal organization in the “resting” steady state of the plasma membrane that is poised to respond to a stimulus. We extended the capabilities of ImFCS by developing a straightforward statistical analysis to provide both spatial maps and highly precise values for diffusion and nanoscale confinement of membrane constituents. We applied these measurements to a panel of structurally distinctive probes (Figure 1B): Using fluorescent protein constructs and selected membrane anchors we targeted Lo-like (palmitoyl-myristoyl (PM) and glycosylphosphatidylinositol (GPI)) and Ld-like (geranylgeranyl (GG)) lipid environments in inner and outer leaflets of the plasma membrane. We also examined the diffusion properties of Lyn kinase, a 40 kDa protein that is anchored to the inner leaflet by PM chains, and IgE-FcεRI receptor complex, which we compared with other transmembrane proteins. Leveraging the unprecedented statistics offered by ImFCS we are able to distinguish characteristic populations of diffusants for each of the probes tested, reflecting the membrane regions through which they travel. With a primary focus on the inner leaflet probes, our data provide strong evidence for previous indications that Lo-like regions are the major component of RBL plasma membranes in the resting steady-state. We also demonstrate that filamentous actin regulates the membrane organization as reflected by changes in the diffusion properties of membrane constituents.

## RESULTS

### Diffusion of probes in the plasma membrane exhibits spatial and temporal heterogeneity – EGFP-GG diffusion in the inner leaflet as an illustrative example

We employed a panel of probes to assess the phase-like membrane heterogeneity in resting RBL cells (Figure 1B). We focus initially on lipid-anchored probes that dynamically partition differentially into Lo-like proteolipid nanodomains as defined in the context of the Kusumi model in the Introduction. These nanodomains exist within the inner and outer leaflets of the plasma membrane and are surrounded by regions of Ld-preferring lipids and proteins; protein complexes may exist in both regions. We posit that diffusion of lipid-anchored probes is relatively faster in Ld-like membrane environments and retarded by their dynamic partitioning into the proteolipid nanodomains, thereby undergoing cycles of free and retarded diffusion (14, 17, 29) (Supplemental Appendix). If a lipid-anchored probe also contains protein modules, these modules may contribute additionally to the level of confinement by the nanodomains as well as to protein-based interactions outside. The case of transmembrane proteins is more complicated because these may have a variety of protein interactions in both leaflets of the membrane (6), and may be surrounded by a lipid “shell” that may affect diffusion and partitioning (54–56).

To evaluate subtle interactions that influence diffusion properties for each probe, we take advantage of the robust statistics and precise quantification offered by ImFCS. The spatial resolution of our current measurements is 2×2 pixels (320×320 nm^2^ = one Px unit; Figure 1A and Supplemental Figure S1A). The *D* value obtained for each Px unit (Equation 1) averages over diffusion within and through the membrane delineated by the actin meshwork (Figure 1A). Compilation from all Px units across the membrane surface provides a distribution of *D* values reflecting the range of membrane environments experienced by a diffusing probe. In our model slower values of *D* represent Px units that are richer in Lo-like proteolipid nanodomains into which the probe partitions to some extent (Supplemental Appendix). FCS is sensitive only to mobile probes, and we examined probes reported to be greater than 85% mobile in non-stimulated cells (45, 57, 58).

We illustrate our experimental analysis with enhanced green fluorescent protein (EGFP) tagged to a short protein sequence that includes a polybasic motif and an acylation site for unsaturated geranyl-geranyl (GG) lipid anchor (EGFP-GG), causing its localization to the membrane inner leaflet (59). EGFP-GG is known to prefer Ld-like regions of the plasma membrane, and partitions relatively weakly into Lo-like regions (59). Figure S1B shows fitting of raw ACFs for each Px unit across the ventral surface (ROI ~8×8 μm^2^) of a representative RBL cell expressing EGFP-GG, from which a spatial *D* map is generated. Although a Brownian diffusion model provides a good fit at the Px unit scale, the distribution of *D* values obtained for EGFP-GG suggests variable area covered by Lo-like nanodomains within Px units.

The presence of Lo-like nanodomains and other elements that retard diffusion is further quantified with svFCS analysis carried out on the same data set for each probe (EGFP-GG in this illustration, Figure S1C). A grouping of 8×8 pixels (1.28×1.28 μm^2^ = Sv unit) is sufficient to create four observation areas of variable sizes (2×2–5×5 pixels) by integrating the fluorescence signal from adjoining pixels in each size group and correlating the fluctuations (60). Resulting ACFs are fitted to obtain a single effective diffusion time *τ*_*D*_ for each observation area size (*A*_eff_) and the four points, *τ*_*D*_ vs *A*_*eff*_, are fitted with a linear model (Equation 2). The constant *slope* is the inverse effective diffusion coefficient (1/*D*_*eff*_), which is proportional to the apparent viscosity experienced by this probe on the micrometer length-scale of the Sv unit (52, 61). Nanoscale information comes from extrapolation of *τ*_*D*_ vs *A*_*eff*_ to *A*_*eff*_ = 0 yielding a *τ*_*0*_ value for each Sv unit. The value of *τ*_*0*_ is expected to be zero for a freely diffusing probe, and the degree to which *τ*_*0*_ is greater than zero provides a relative measure of confinement of that probe in domains on nanoscale smaller than a Px unit. Repeating svFCS analysis over all possible non-overlapping Sv units yields a *τ*_*0*_ map and a *Slope* = I/*D*_*eff*_ map for that cell (Figures S1C, inset and S2).

We utilize the same ImFCS data set to evaluate temporal heterogeneity in the spatially resolved *D* (Px units) and *τ*_*0*_ and *Slope* (Sv units) maps in shorter time windows by dividing the entire raw image series (80,000 frames, 3.5 ms/frame; Figure S2A) into four equal 70 seconds segments and conducting ACF and svFCS analyses on each segment. As shown in Figure S2, the *D*, *τ*_*0*_, and *Slope* maps for EGFP-GG show moderate temporal fluctuations across the measured ventral membrane, i.e., the values of each of these parameters for any given region change somewhat across the time segments. However, there are no obvious regions of the maps that look distinctively different at this level of resolution, and the distributions of *D*, *τ*_*0*_, and *Slope* values across all pixels for each segment remain very similar (Figure S2B). Therefore, we take the spatial heterogeneity of *D*, *τ*_*0*_, and *Slope* values to represent the distribution of these diffusion properties exhibited by the probe as it explores the plasma membrane. Similar to EGFP-GG, we observed local fluctuations around overall averages (Figure S2) for all plasma membrane probes we evaluated in this study (Figure 1B), underscoring the dynamics of the plasma membrane in the resting steady-state of RBL cells. We further observed for all probes that the cell-averaged values of *D*, *τ*_*0*_, and *Slope* remain roughly the same from cell to cell (evaluated on different days), allowing us to pool unit values across the spatial maps of multiple cells. The *D*, *τ*_*0*_, and *Slope* values quantify the diffusion properties of a given probe without providing specific information about the physico-chemical interactions involved (e.g., lipid- or protein-based). However, the nature of these interactions and the heterogeneity of the plasma membrane can be inferred by comparing the diffusion properties of structurally different probes, which experience the same membrane differentially.

Representative examples of *D* values for 18 cells expressing EGFP-GG are shown in Figure 2A. The measured ROI for each cell surface includes about 625 Px units, such that pooling all *D* values for these many cells gives a very large number (>10,000 *D* values) for EGFP-GG (or another probe) that is not biased from a subset of cells. The statistically robust arithmetic average, *D*_*av*_ = 0.64 ± 0.002 μm^2^/s (mean ± SEM) we obtain for EGFP-GG (Table 1) is comparable to those obtained from other types of diffusion measurements (62, 63). More detailed information is obtained from compiling the pooled data as a cumulative distribution function (CDF; Figure 2B, dashed line). These same data may also be plotted as a probability distribution function (PDF; Figure 2B, inset), which allows easier visualization of underlying Gaussian components. CDFs, which are mathematically equivalent to PDFs but do not require arbitrary range-binning of parameter values, are particularly useful for statistical analyses. We fitted the CDFs of *D* values with either one- or two-component Gaussian models (Equations 3 and 4), and determined the best fit by comparing residuals and reduced chi-squared values (Figures 2B and S3). Values of *τ*_*0*_ and *Slope* compiled over many cells can also be plotted as a CDF or PDF and fitted. However, the statistics are not as robust compared as the *D* values (~36 Sv units/cell, ~500 for ~15 cells), and we use the arithmetic averages *τ*_*0,av*_ and *Slope*_*av*_ to compare probes.

**Figure 2.**
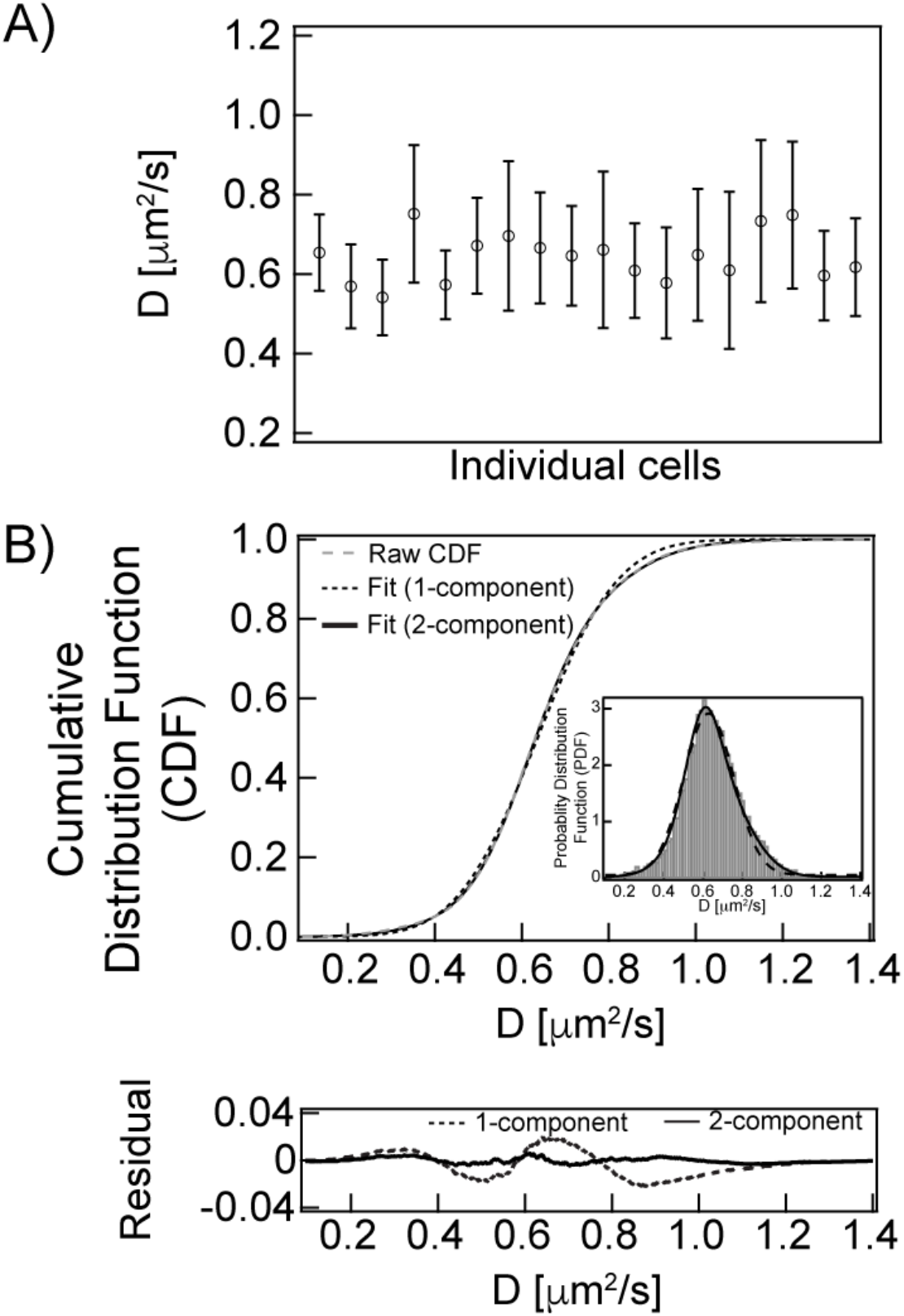
Very large data sets from ImFCS measurements are analyzed to examine spatially heterogeneous diffusion properties of plasma membrane probes, as exemplified by EGFP-GG. **A**) *D* values averaged over ROIs in 18 individual RBL cells expressing EGFP-GG; error bars are standard deviations about an average *D* for each cell. ROIs generally contain 400-625 Px units covering about 41-64 μm2 membrane area for each cell, yielding 10,527 total *D* values for 18 cells in this example. **B**) Top: *D* values obtained from ROIs in all cells are pooled and plotted as a normalized, cumulative distribution function (CDF), which is fitted with one or two components, as indicated. The inset shows the same data for *D* plotted as a probability distribution function (PDF) with arbitrary binning of parameter values. Bottom: Residual plots for one-component (Equation 3) and two-component (Equation 4) fits to CDF.

**Table 1.**
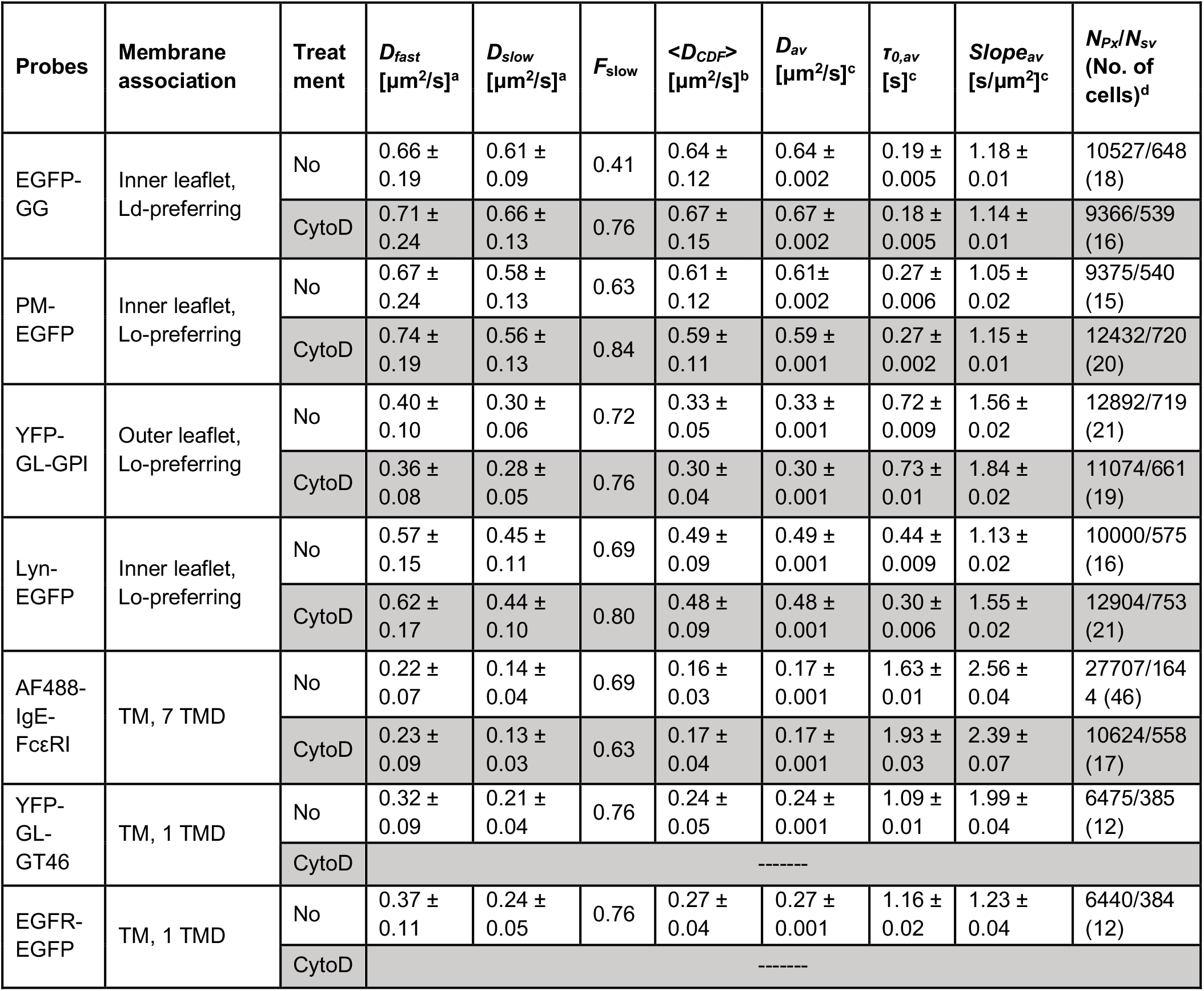
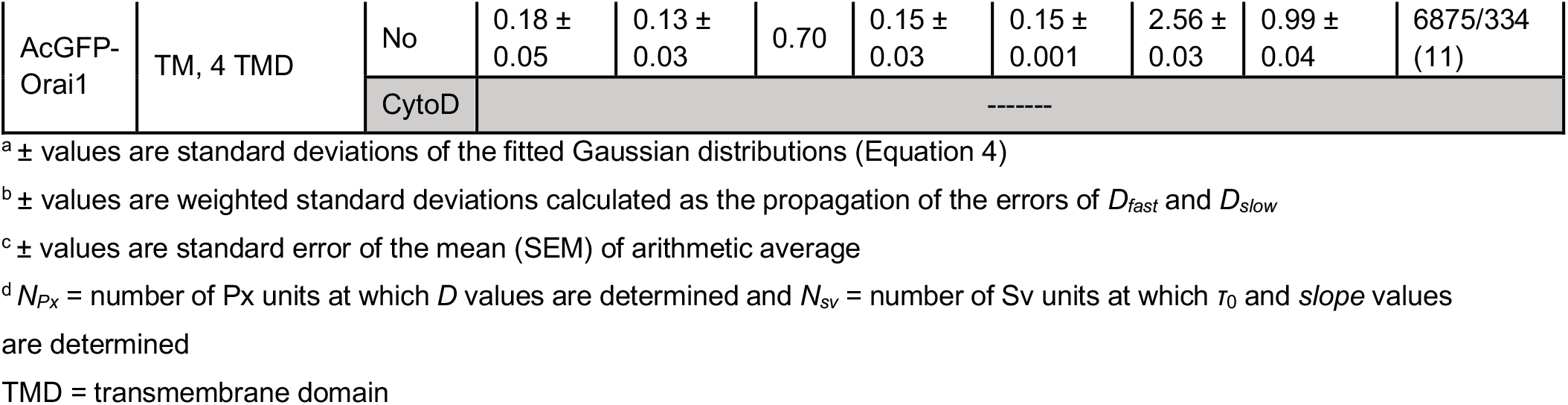
Fitting results of experimental CDFs of *D;* and *τ*_*0,av*_ and *slope*_*av*_ of membrane probes shown in Figure 1, without and with CytoD treatment.

The CDF of *D* values for EGFP-GG in Figure 2B is not fit satisfactorily with a single Gaussian population of diffusing probes (Equation 3), consistent with some partitioning of EGFP-GG into nanodomains. A good fit requires at least two Gaussian components (Equation 4), which we interpret as differences in nanodomain coverage among Px units that cause this probe’s diffusion properties to group into two populations. Each of these distinguishable populations of Px units may represent averages over sub-populations. The fractional amounts of each component, *F*_*slow*_ and *F*_*fast*_ = 1 − *F*_*slow*_, represent the fraction of Px units that are nanodomain-rich (*D*_*slow*_) or nanodomain-poor (*D*_*fast*_), respectively, as experienced by a given probe depending on its partitioning into nanodomains. Although distinguishable, the values of the two *D* components do not differ by much for EGFP-GG (Table 1, Figure 3): *D*_*slow*_ = 0.61 ± 0.09 μm^2^/s (*F*_*slow*_ = 0.41) and *D*_*fast*_ = 0.66 ± 0.19 μm^2^/s (*F*_*fast*_ = 0.59), where the ± values are standard deviations (*σ*) of the fitted Gaussian distributions and represent the width of the respective diffusing population (Figure 2B; Equation 4). As we demonstrate in the Supplement (Table S1, Figure S3), very large pooled data sets (*N*_*Px*_ ~10,000; Table 1) allow distinctions as small as 10% to be made for *D*_*fast*_ and *D*_*slow*_ components. The weighted average (Equation 5), *F*_*fast*_ *D*_*fast*_ + *F*_*slow*_ *D*_*slow*_ = *D*_*CDF*_ = 0.64 ± 0.12 μm^2^/s, which is the same as *D*_*av*_, the arithmetic average of the total pooled *D* values for EGFP-GG.

**Figure 3.**
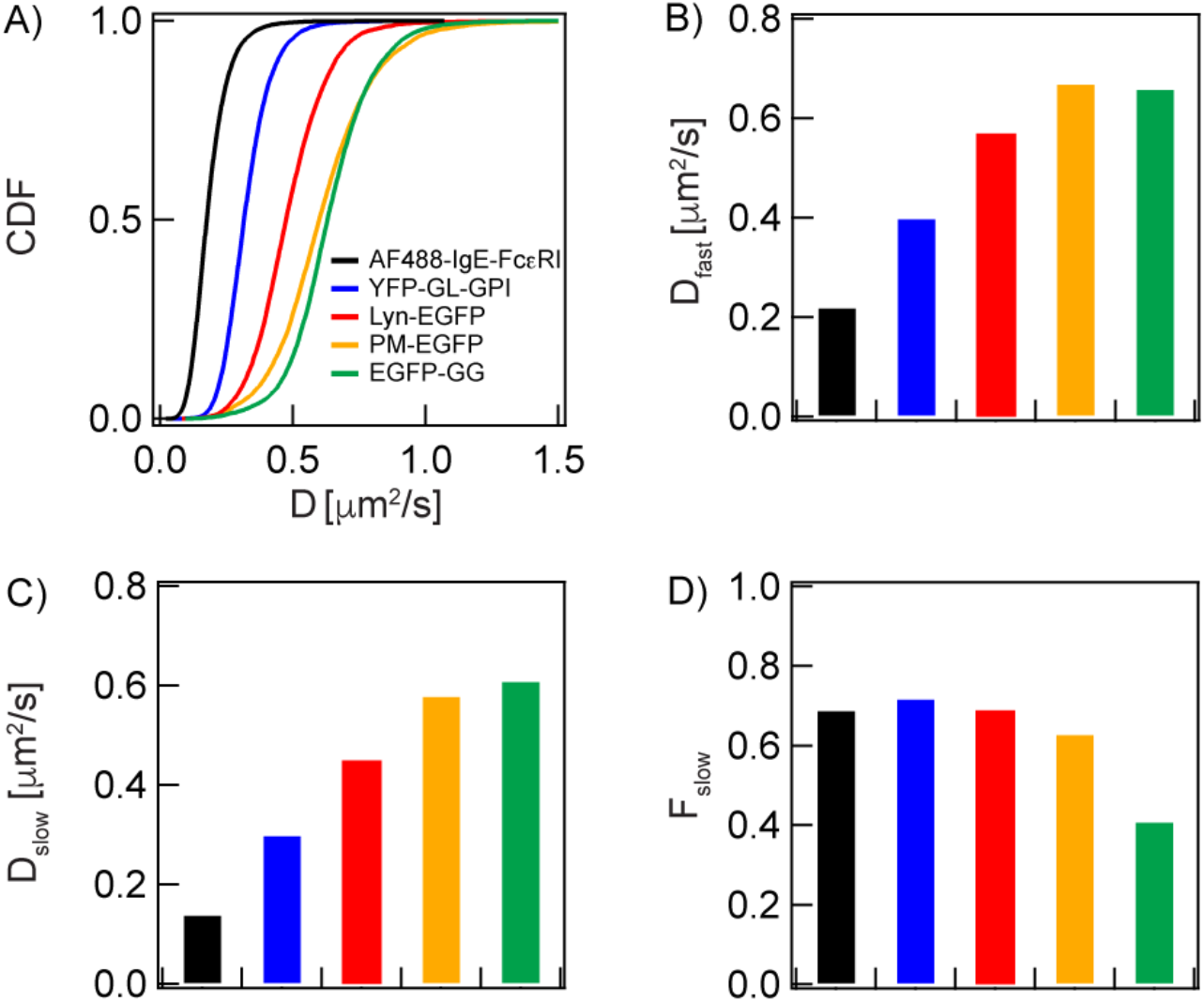
Diffusion parameters are determined from the statistical analyses of *D* CDF for lipid-anchored and TM probes depicted in Figure 1B. **A**) CDF of *D* values for the probes, as indicated. Fitting of the respective CDFs yields: **B**) *D*_fast_: diffusion coefficient for Px unit population with less dynamic confinement (nanodomain-poor for lipid-anchored probes); **C**) *D*_slow_: diffusion coefficient for Px unit population with more dynamic confinement (nanodomain-rich for lipid-anchored probes); **D**) *F*_slow_: Fraction of Px units with the population exhibiting *D*_slow_. The lipid-anchored probes (EGFP-GG, PM-EGFP, Lyn-EGFP, YFP-GL-GPI) are primarily subject to lipid-based interactions whereas the TM probe (AF488-IgE-FcεRI) depends on protein-based interactions. The color code in (A) identifies the probes in all panels. Numerical values of all parameters with defined errors are provided in Table 1.

To explore the spatial distribution of slower and faster EGFP-GG diffusers and their connectivity we made contour maps of *D* values on individual cells. Contour maps for one cell divided in 70 s time segments (as described above) confirms the dynamic heterogeneity of the plasma membrane as sensed by this probe (Figure S4A). Contours maps of EGFP-GG *D* values determined from the standard (280 s) data acquisition period and compared for different single cells show diversity in distributions and connectivity, although the *D*_*av*_ values for each cell are very similar (Figures 2 and S4B).

The averaged *τ*_*0*_ value for EGFP-GG (*τ*_*0,av*_ = 0.19 ± 0.12 s) falls just within the range that cannot be confidently distinguished from zero experimentally (0<*τ*_*0*_<0.2 s; (64)), reflecting relatively little retardation by nanodomains. Together, the *D*_*fast*_, *D*_*slow*_, and *τ*_*0,av*_ values for EGFP-GG reveal the presence of nanodomains that are detectable by our measurements. *F*_*fast*_ and *D*fast represent the larger of two populations of Px units that are sensed as nanodomain-poor, further consistent with EGFP-GG partitioning relatively weakly into nanodomains.

#### EGFP-GG, PM-EGFP, and Lyn-EGFP diffuse differentially in the membrane inner leaflet

Lyn kinase, which is anchored to the inner leaflet of the plasma membrane by saturated palmitoyl (P) and myristoyl (M) chains, is involved in the earliest stage of transmembrane signaling triggered by antigen-crosslinking of IgE-FcεRI (Introduction). PM-EGFP, which is constructed from the small segment of Lyn that is acylated, has been established as an inner leaflet probe that partitions preferentially into Lo-like environments of the plasma membrane, in contrast to EGFP-GG (59, 65). Lyn has additional cytosolic protein modules, including SH3, SH2, and kinase modules. We directly compared PM-EGFP to EGFP-GG and to Lyn-EGFP to determine how lipid-based and protein-based interactions influence the diffusion properties of inner leaflet probes, including differential confinement in nanodomains (Figure 3).

As averaged over many Px units in multiple cells, *D*_*av*_ for PM-EGFP is 0.61 μm^2^/s, which is within the range of literature reports for similar probes (66–69), and slower than that for EGFP-GG, which partitions relatively weakly into Lo-like environments (Table 1). The CDF of *D* values for PM-EGFP is satisfactorily fitted with two population components, and the larger fraction has the slower diffusion coefficient: *D*_*slow*_ = 0.58 μm^2^/s (*F*_*slow*_ = 0.63) and *D*_*fast*_ = 0.67 μm^2^/s (*F*_*fast*_ = 0.37) (Figure 3, B–D). We consider that PM-EGFP interacts with the plasma membrane by means of its saturated fatty acyl chains and that these chains are slowed in their lateral diffusion by the extent of their interactions with nanodomains. Consistent with their differential preference for Lo-like environments, their respective *F*_*slow*_ values (Table 1, Figure 3D) indicate that diffusing PM-EGFP is more sensitive than EGFP-GG to the presence of nanodomains. Contour maps of *D* values for PM-EGFP and EGFP-GG for individual cells show consistent results that regions of slower diffusion are more pronounced for PM-EGFP compared to EGFP-GG (Figure S4B). Moreover, the *F*_*slow,cell*_ values determined from contour maps of individual cells show significantly higher values for PM-EGFP compared to EGFP-GG, consisted with differences determined from fitting CDF of data ensemble (Figure S4C; Table 1)

The *τ*_*0,av*_ of PM-EGFP (0.27 s) is larger than that for EGFP-GG (0.19 s) (Table 1, Figure 4E), similarly corresponding to greater confinement of PM-EGFP in nanodomains. The results for both PM-EGFP and EGFP-GG can be explained by a membrane model with nanodomain-rich and nanodomain-poor regions (*F*_*slow*_, *F*_*fast*_), and differences in *F*, *D*, and *τ*_*0*_ values reflect the degree of nanodomain confinement experienced by a particular probe (Appendix, Scheme A3). The value of *Slope*_*av*_ for PM-EGFP (1.05 s/μm^2^) compared to that for EGFP-GG (1.18 s/μm^2^) indicates that the apparent micrometer-scale viscosity is greater for EGFP-GG, suggesting that this probe is more slowed than PM-EGFP by interactions outside nanodomains.

**Figure 4.**
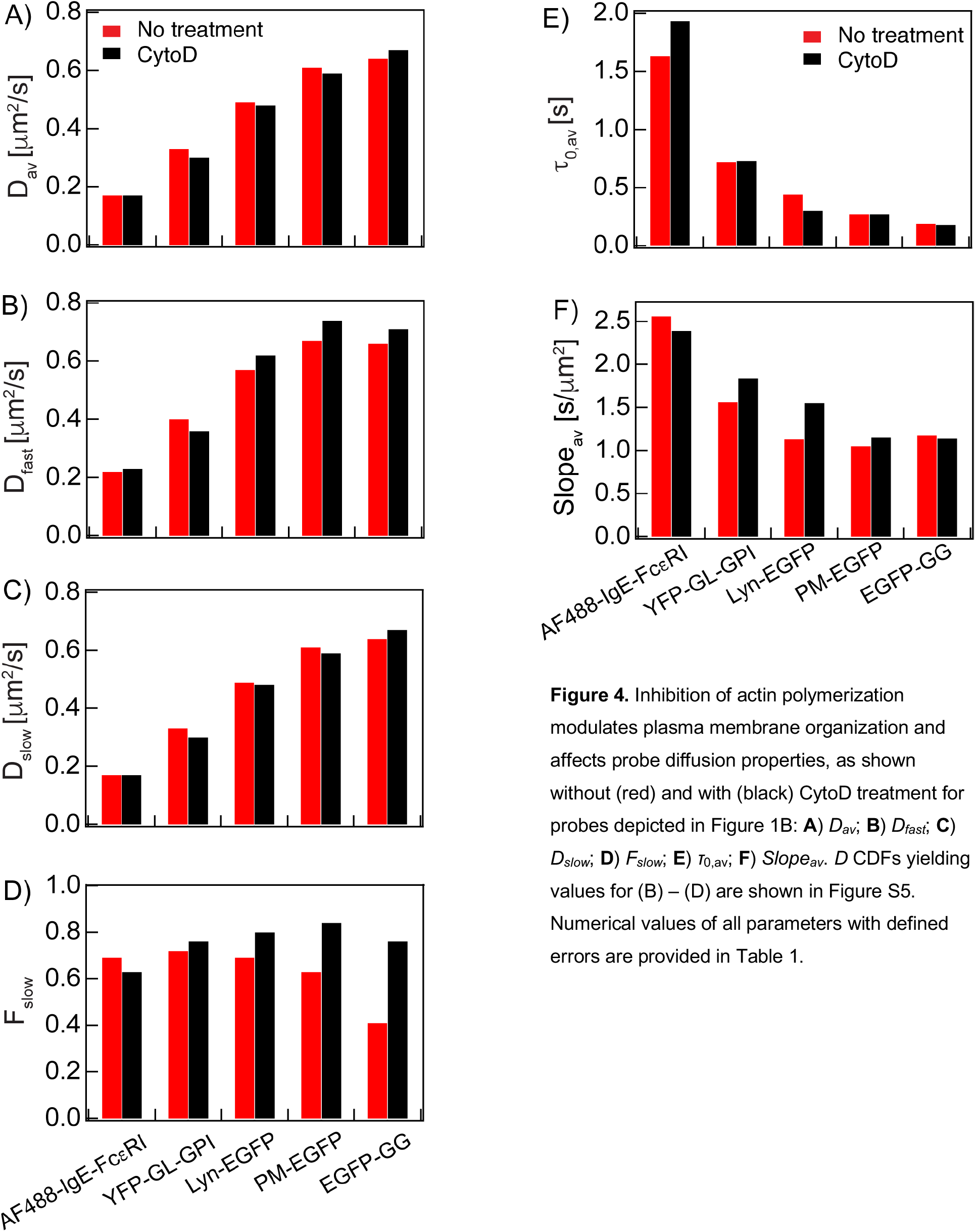
Inhibition of actin polymerization modulates plasma membrane organization and affects probe diffusion properties, as shown without (red) and with (black) CytoD treatment for probes depicted in Figure 1B: **A**) *D*_*av*_; **B**) *D*_*fast*_; **C**) *D*_*slow*_; **D**) *F*_*slow*_; **E**) *τ*0,av; **F**) *Slope*_*av*_. *D* CDFs yielding values for (B) – (D) are shown in Figure S5. Numerical values of all parameters with defined errors are provided in Table 1.

The *D*_*av*_ for Lyn-EGFP (0.49 μm^2^/s) is markedly slower than that for PM-EGFP (Table 1) and within the range reported previously for Lyn and other src family kinases (68–70). The CDF of *D* values for Lyn-EGFP resolves into two populations, with the larger fraction of Px units experienced by this probe as nanodomain-rich: *D*_*slow*_ = 0.45 μm^2^/s (*F*_*slow*_ = 0.69); *D*_*fast*_ = 0.57 μm^2^/s (Figure 3, Table 1). These values, compared to those of PM-EGFP, are consistent with Lyn-EGFP interacting more strongly with nanodomains such that this probe diffuses more slowly in both nanodomain-rich and nanodomain-poor Px units (Appendix, Scheme A3). The substantially larger *τ*_*0,av*_ value for Lyn-EGFP (0.46 s) compared to those for PM-EGFP and EGFP-GG (Table 1, Figure 4E) further indicates that Lyn-EGFP has more interactions to increase confinement on the nanoscale. The *Slope*_*av*_ for Lyn-EGFP (1.13 s/μm^2^) is greater than that for PM-EGFP and less than that EGFP-GG (Table 1; Figure 4F). Given the clear differences in diffusional properties and assuming the saturated PM acyl chains for both PM-EGFP and Lyn-EGFP are similarly restricted by Lo-like nanodomains, the cytosolic protein modules of Lyn-EGFP appear to be interacting with additional sources of confinement within nanodomains and possibly with separate protein complexes. As described in a subsequent section, we used inhibition of actin polymerization to further distinguish these contributions.

### Lipid-anchored probe in the outer leaflet diffuses differently from those in the inner leaflet

We evaluated YFP-GL-GPI, a glycosylphosphatidylinositol (GPI) anchored protein, as an outer leaflet probe that partitions favorably into Lo-like environments (71). We find *D*_*av*_ = 0.33 μm^2^/s (Table 1), which is within the range of values previously reported for this probe (45, 63, 67, 72, 73). Notably, *D*_*av*_ is markedly slower and *τ*_*0,av*_ (0.72 s) is markedly higher than inner leaflet probes PM-EGFP and Lyn-EGFP (Table 1; Figure 4, A and E). These observations point to substantive differences in the physical properties that affect diffusion in the inner *vs* outer leaflet, particularly as related to confining nanodomains. The *D* CDF for YFP-GL-GPI resolves into two populations of Px units with the larger population showing the slower diffusion coefficient for this probe (Table 1; Figure 3): *D*_*slow*_ = 0.30 μm^2^/s (*F*_*slow*_ = 0.72) and *D*_*fast*_ = 0.40 μm^2^/s. The fractional amounts indicate that YFP-GL-GPI, similar to PM-EGFP and Lyn-EGFP, exhibits slower diffusion in the bulk (60-70%) of the membrane sensed as nanodomain-rich Px units. Consistent with the slower *D* and higher *τ*_*0,av*_, the *Slope*_*av*_ value for YFP-GL-GPI is larger compared to inner-leaflet Lo-preferring probes PM-EGFP and Lyn-EGFP (Table 1; Figure 3F), reflecting differences in the interactions that retard diffusion for each probe inside and outside of nanodomains.

#### Inhibition of actin polymerization differentially affects diffusion of Lyn-EGFP and other lipid anchored probes

The dynamic actin cytoskeleton has been shown to interact, directly or indirectly, with membrane localized proteins, affecting their functions (74, 75). Therefore, to gain insight into additional interactions of Lyn-EGFP compared to its lipid anchor alone (PM-EGFP), we evaluated effects of cytochalasin D (CytoD), which acutely inhibits actin polymerization (Figures 4 and S5; Table 1). We found that treatment of RBL cells with 1 μM CytoD causes *Dav* values to decrease for both Lyn-EGFP and PM-EGFP, with similar trends in *D*_*slow*_ (slightly decrease) and *D*_*fast*_ (increase) (Figures 4, A–C and S5A; Table 1). *F*slow increases, indicating more Px units are sensed as nanodomain-rich by both probes (Figure 4D). Changes in their respective *τ*_*0,av*_ and *Slope*_*av*_ values after CytoD treatment clearly differentiate Lyn-EGFP from PM-EGFP. Whereas *τ*_*0,av*_ decreases from 0.46 to 0.31 s and *Slope*_*av*_ increases from 1.13 to 1.55 s/μm^2^ for Lyn-EGFP, the value of these parameters stay about the same (*τ*_*0,av*_) or increase much less (*Slope*_*av*_) for PM-EGFP (Figures 4E,F and S4B,C; Table 1). Thus it appears that values observed for Lyn-EGFP in untreated cells depend in part on protein-mediated interactions, which in turn depend on cytoskeletal organization that is perturbed by inhibition of actin polymerization. Notably, the values for *τ*_*0,av*_ for PM-EGFP (0.27 s) and Lyn-EGFP (0.31 s) in CytoD treated cells are similar, suggesting that the two probes are confined by nanodomains similarly under these conditions, as driven largely by their lipid anchors. The increased *Slope*av for Lyn-EGFP indicates that CytoD treatment increases this probe’s protein-based interactions outside nanodomains, resulting in an increase in apparent membrane viscosity on micrometer lengthscale.

The *D*, *τ*_*0,av*_, and *Slope*_*av*_ values for EGFP-GG diffusing in membranes of cells, without and with CytoD treatment provide additional information about changes occurring in the membrane inner leaflet (Table 1). Whereas treatment with CytoD causes *D*_*av*_ values to decrease for Lo-preferring Lyn-EGFP and PM-EGFP, this value increases for Ld-preferring EGFP-GG (Figures 4A and S5A). For EGFP-GG, *D*_*slow*_ with treatment is the same as *D*_*fast*_ without treatment (0.66 μm^2^/s) and *D*_*fast*_ becomes even faster (0.71 μm^2^/s) with treatment, while *F*_*slow*_ increases from 0.41 to 0.76 (Figure 4, B-D). These changes suggest that EGFP-GG partitions even less into nanodomains after treatment, thereby diffusing faster in Px units sensed as nanodomain-poor, even as the fraction of Px units sensed as nanodomain-rich (*F*_*slow*_) increases. Similar to PM-EGFP, values for *τ*_*0,av*_ remain about the same for EGFP-GG before and after CytoD treatment (Figures 4E and S5B), but the value for *Slope*_*av*_ decreases by a small amount, rather than increasing as for PM-EGFP and Lyn-EGFP (Figures 4C and S5C). Collectively, our results indicate that CytoD treatment causes Lo-like nanodomains to become more ordered and more Px units to become relatively more nanodomain-rich in the membrane inner leaflet, and also that this treatment alters Lyn-EGFP’s protein-based interactions inside and outside of nanodomains.

Although YFP-GL-GPI in the outer leaflet diffuses more slowly than PM-EGFP in the inner leaflet, CytoD treatment causes *D*_*av*_ to decrease and *F*_*slow*_ to increase for of all of the Lo-preferring probes (Figures 4A–D and S5A; Table 1), suggesting the ordered lipid character of nanodomains and their coverage area increases in both leaflets of the plasma membrane. The *τ*_*0,av*_ value is similarly unchanged for both YFP-GL-GPI and PM-EGFP after treatment (Figure S5B), indicating that the net level of confinement remains similar. However, *Slope*_*av*_ increases markedly for YFP-GL-GPI (Figure S5B), pointing to additional interactions inside and outside of nanodomains such that the apparent membrane viscosity on the micrometer scale increases after treatment as experienced by this probe.

### AF488-IgE-FcεRI and other transmembrane proteins exhibit additionally restricted diffusion

TM proteins have additional potential for interacting with other proteins, and various types and strengths of these interactions (specific, nonspecific, steric) are likely to be complex. In most cases, protein-based interactions probably dominate over tendencies to partition into Lo-like nanodomains although palmitoylation, for example, may modulate these interactions (76). We monitor the TM receptor FcεRI as a complex with Alexafluor 488 labeled IgE (AF488-IgE-FcεRI; Figure 1B). As quantified with *Dav* (0.17 μm^2^/s) and *τ*_*0,av*_ (1.65 s), FcεRI diffuses more slowly on the scale of Px units and is more highly confined nanoscopically than Lyn-EGFP and all the inner and outer leaflet, lipid-anchored probes (Table 1). This *D*_*av*_ value agrees well with previous reports (77–79). The *D* CDF of AF488-IgE-FcεRI is fitted with two population components: *D*_*slow*_ = 0.14 μm^2^/s (*F*_*slow*_ = 0.69) and *D*_*fast*_ = 0.22 μm^2^/s (Figure 3). We observed small but significant changes in these values after treatment with CytoD (Table 1; Figures 4 and S5)

We also evaluated three other TM protein probes in resting RBL cells: epidermal growth factor receptor (EGFR-EGFP (80)), GT46 (YFP-GL-GT46 (71)), and Ca^2+^ channel Orai1 (AcGFP-Orai1 (81)). In monomeric form, EGFR and GT46 (TM segment of LDL receptor and cytoplasmic tail of CD46) have a single TM segment, whereas Orai1 has four and FcεRI has seven TM segments. The CDFs of *D*, *τ*_*0,av*_, and *Slope*_*av*_ values for all four TM probes are shown with key parameters summarized in Table 1 and Figure S6. As expected, the *D*_*av*_ of these protein probes are consistently slower than the lipid-anchored probes, and those with a single TM segment diffuse somewhat faster: YFP-GT46 (0.26 μm^2^/s) and EGFR-EGFP (0.24 μm^2^/s), compared to AcGFP-Orai1 (0.15 μm^2^/s) and AF488-IgE-FcεRI (0.17 μm^2^/s). We also observed bimodal *D* CDFs with *F*_*slow*_ = 0.70-0.76 for these other TM probes. Thus, all TM protein probes tested distributed detectably into two diffusing populations, reflecting differences in membrane environments at the spatial resolution of Px units.

All the protein probes show substantial nanoscale confinement as represented by relatively high *τ*_*0,av*_ values: 1.14 s (YFP-GT46), 1.41 s (EGFR-EGFP), 1.63 s (AF488-IgE-FcεRI), and 2.56 s (AcGFP-Orai1) (Table 1; Figure S5). There are multiple possible sources for confinement of each of these TM probes, including protein-based interactions with cellular constituents proximal to and within the plasma membrane. The *Slope*_*av*_ values show no obvious trends related to number of TM segments: among these four probes AF488-IgE-FcεRI (2.56 s/μm^2^) experiences the highest apparent viscosity on the micrometer scale, AcGFP-Orai1 (0.99 s/μm^2^) the lowest, and those with a single TM segment in between. Although the particular contributions of protein-based interactions cannot be discerned by comparing these probes, our results confirm expectations that proteins with TM segments are more restricted in diffusion compared to lipid-anchored membrane probes and provide quantitative details of their distinctive diffusion properties.

## DISCUSSION

Our study demonstrated the versatility and quantitative rigor of ImFCS to examine RBL mast cells with two primary purposes: 1) characterize the dynamic heterogeneity of the resting, poised plasma membrane as sensed by a panel of structurally distinct probes; 2) establish a foundation for elucidating subtle changes that occur when this poised plasma membrane responds to a stimulus and initiates transmembrane signaling. We are motivated by IgE-FcεRI coupling with lipid-anchored Lyn kinase in Lo-like proteolipid nanodomains that are stabilized after antigen stimulation (23).

We evaluated lipid-anchored probes in the membrane inner and outer leaflets and TM protein probes in resting RBL cells, and obtained spatial maps of *D* (Px units), *τ*_*0*_, and *Slope* (Sv units) values for each cell (Figures 1 and S1). Similar values of these parameters for a given probe across cells and over time (Figures 2A and S2) allowed us to combine data from multiple cells (~10,000 Px units; ~500 Sv units), yielding exceptionally robust quantification for specified diffusion properties. The precision of these values, e.g., very small SEM of the arithmetic averages, sharpens the contrast between different probes diffusing through the same membrane milieu (Table 1). The data-intense CDFs of *D* values are fitted (Equation 4) to identify as small as 10% differences in lateral diffusion (*D*_*fast*_ vs *D*_*slow*_) of a given probe, which arise from variable degrees of weak nanoscopic interactions within different Px units (Figure S3). Corresponding to *D*_*fast*_ and *D*_*slow*_, we define these regions as nanodomain-poor and nanodomain-rich, respectively. *F*_*slow*_ is the fraction of Px units sensed as nanodomain-poor. The diffusion-law equation (Equation 2) applied to Sv units provides indirect information about a probe’s diffusion properties at both nanoscopic (*τ*_*0*_) and microscopic (*Slope =* 1/*D*_*eff*_) length scales.

We primarily examined lipid-anchored probes with contrasting acyl chains that prefer Lo-like or Ld-like environments, and we interpret our data in terms of a simple model that considers Lo-like proteolipid nanodomains within regions of Ld-preferring lipids and proteins (Figure 1; Appendix). This view fits within the hierarchal model for plasma membrane organization as proposed by Kusumi and colleagues and extended in other laboratories (Introduction). The distinction between nanodomain-rich and nanodomain-poor Px units may be, for example, respectively higher and lower densities of cortical actin meshwork in the ‘hierarchal model’ (24) or of acto-myosin asters in the ‘active composite model’ (38). Contour maps show the connectivity of slower and faster *D* values as they differ for Lo-preferring and Ld-preferring probes at Px unit resolution (Figure S4).

### Inner leaflet probes sense the membrane differentially

We integrate the diffusion behaviors of EGFP-GG, PM-EGFP and Lyn-EGFP into a common inner leaflet model (Figure 5A). A larger *Dav* value and smaller *τ*_*0,av*_ value for EGFP-GG compared to the other two lipid-anchored probes is consistent with the view that the unsaturated geranyl-geranyl acyl chains are less dynamically confined in Lo-like nanodomains than are saturated palmitate-myristoylate chains. Correspondingly, a lower fraction of Px units have sufficient nanodomains to yield the slower diffusing population (*F*_*slow*_, *D*_*slow*_) for EGFP-GG compared to PM-EGFP (Figure 3B,D,E). Precise numerical details in Table 1 provide additional insight to these subtle interactions. For example, values for *D*_*fast*_ and *D*_*slow*_ are not very different for these two probes and are most similar to each other for *D*_*fast*_, as might be expected when both probes are diffusing in Ld-like Px units.

**Figure 5:**
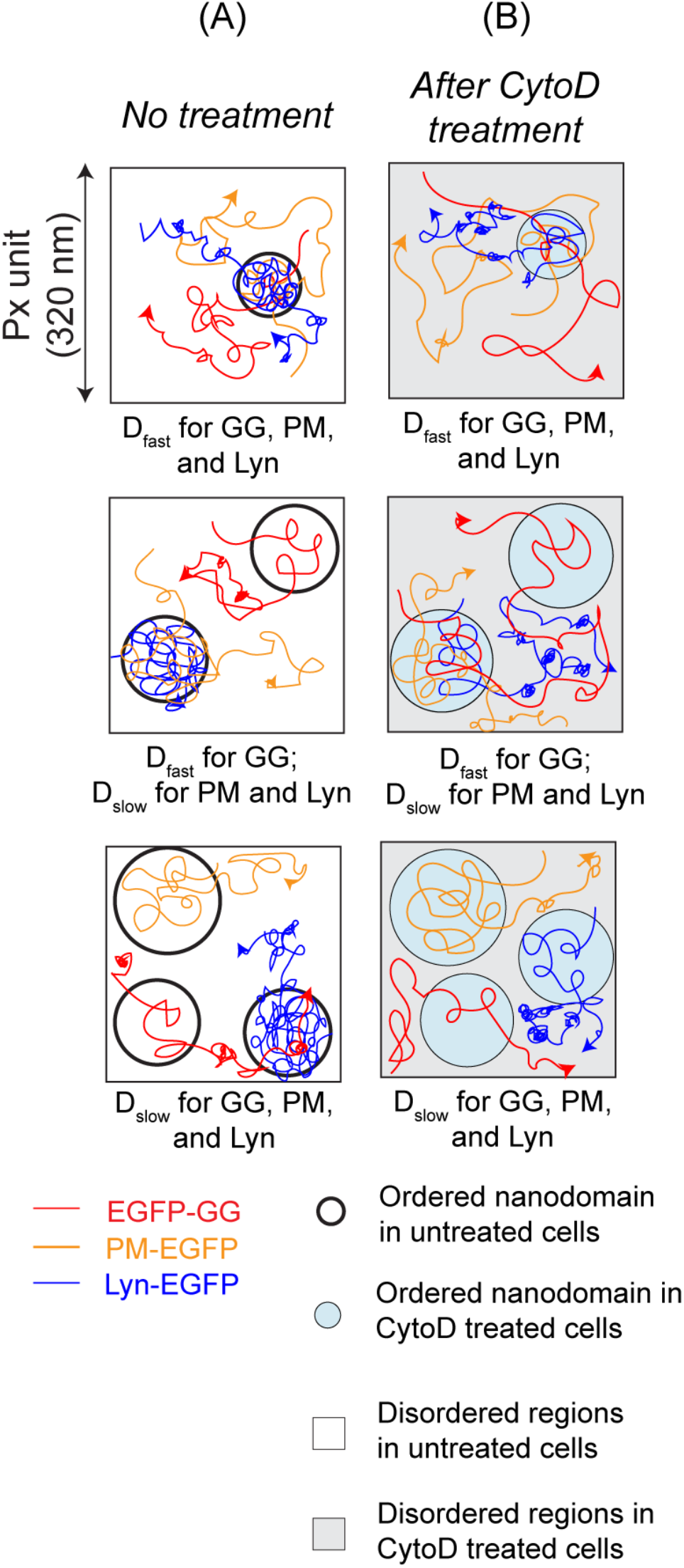
Diffusion of inner leaflet, lipid-anchored probes depends on interactions inside and outside of Lo-like proteolipid nanodomains and is modulated by inhibition of actin polymerization. Representative Px units with low (top) moderate (middle), or high (bottom) coverage by nanodomain (circles) are shown. **A**) In untreated cells, retardation of diffusion due to dynamic partitioning into nanodomains: Lyn-EGFP > PM-EGFP > EGFP-GG, as reflected by values for *τ*_*0,av*_ (Table 1); the level of interactions outside nanodomains is: EGFP-GG ≥ Lyn-EGFP > PM-EGFP, as reflected by values for *Slope*_*av*_. **B**) After CytoD treatment, the nanodomains become more ordered and the membrane regions outside nanodomains also change. In contrast to PM-EGFP, the confinement of Lyn-EGFP in the nanodomains decreases while its interactions outside nanodomains increase; for PM-EGFP and EGFP-GG, the confinement in the nanodomains does not change significantly, whereas there are modest increases or decreases, respectively, in interactions outside nanodomains. Differences in diffusion properties caused CytoD treatment are also reflected in *D* CDFs and extracted *D* values for inner leaflet probes as described in text and listed in Table 1.

Differences between PM-EGFP and Lyn-EGFP, which have the same lipid anchor, suggest that some nanodomains include proteins that interact with Lyn’s protein modules. Lyn-EGFP has slower *D*_*av*_ and *D*_*slow*_ values and a somewhat larger *F*_*slow*_ value (Figure 3B-E). Similarly, Lyn-EGFP’s slower value for *D*_*fast*_ suggests that the population of Px units with less abundant nanodomains retards diffusion for Lyn-EGFP more than for PM-EGFP or that Lyn-EGFP interacts with other proteins in the membrane or cytoplasm when this probe diffuses outside of the nanodomains. Interestingly, *Slope*_*av*_ for Lyn-EGFP is greater than that for PM-EGFP and less than that EGFP-GG, and this measure of micrometer scale viscosity indicates that PM-EGFP has the lowest level of interactions outside of nanodomains. EGFP-GG’s apparent higher level of interactions may be due to the polybasic motif included in this construct to stabilize its membrane localization, e.g., electrostatic interactions with phosphatidyl inositols or other negatively charged phospholipids that localize to Ld-like regions.

Changes in the diffusion properties of these probes after CytoD treatment are consistent with previous observations that plasma membrane heterogeneity is remodeled upon acute inhibition of actin polymerization (Figures 4 and S4; Table 1) (75). Our measurements point to a consistent picture that (1) the relative number of Px units rich in nanodomains increases, (2) the nanodomain coverage a) increases in nanodomain-rich Px units and b) decreases in nanodomain-poor Px units, and (3) overall the nanodomains become more ordered (Figure 5B). Compared to their counterparts in untreated cells, EGFP-GG diffuses faster in both nanodomain-rich and nanodomain-poor Px units, whereas PM-EGFP and Lyn-EGFP diffuse faster in nanodomain poor Px units and slower in nanodomain rich Px units (Figure S4A). These results provide new evidence that CytoD treated cells exhibit stronger phase-like segregation such that the diffusion behavior of EGFP-GG and PM-EGFP becomes more distinguishable. Previous studies showed that the long chain actin meshwork serves to restrict phase separation (30–34), and that CytoD increases the size of corrals formed by this meshwork (82). Experiments and Ising-based simulations showed that increasing the dimensions of this meshwork increases co-clustering of Lo-preferring components (75). In these simulations, this effect is observed whether Ld-like or Lo-like components (but not both randomly) are pinned to the meshwork. However, the tendency of Lo-like components to co-cluster within the corrals increases monotonically only in the case where Lo-like components are pinned to the meshwork, which may be explained by release of sequestered Lo-like components as the meshwork becomes less dense. Prevention of small, acto-myosin mediated nanodomains may additionally release Lo-like components and increase the tendency for phase-like separation.

CytoD differentially affects the diffusion of Lyn-EGFP compared to PM-EGFP, reflecting Lyn’s protein modules. In contrast to PM-EGFP and EGFP-GG, large changes for Lyn-EGFP in *τ*_*0,av*_ (decrease) and *Slope*_*av*_ (increase) after CytoD treatment (Figures 4 and S4) indicate that Lyn-EGFP’s protein-based interactions decrease within nanodomains and increase outside of nanodomains. Some of these changes may be related to concurrent modulation of Lo-like nanodomains. Lyn may also interact with other partners via its SH2, SH3, and kinase protein modules in an actin-dependent manner. As shown previously for CytoD treated cells, TM proteins tend to cluster in Ld-like regions of the plasma membrane in intact cells (83), and the average size of protein assemblies in plasma membrane sheets increases significantly (84). These changes could retard the diffusion of Lyn-EGFP anchored to the inner leaflet as well as the diffusion YFP-GL-GPI anchored to the outer leaflet and TM probes. Correspondingly, the values of *Slope*_*av*_ go up markedly for all of these probes after CytoD treatment while changing much less for PM-EGFP (increase) and EGFP-GG (decrease).

### Diffusion of Lo-preferring probes differs in outer vs inner leaflets

Lipid-anchored probes PM-EGFP and YFP-GL-GPI both prefer Lo-like environments, and we found that the two leaflets exhibit a similar predominance of nanodomain-rich Px units as sensed by both of these probes with *F*_*slow*_ values of 0.63 and 0.72, respectively (Figure 3E, Table 1). However, absolute values of *D*_*slow*_, *D*_*fast*_, and *D*_*av*_ of YFP-GL-GPI are substantially slower than those values for PM-EGFP, indicating greater sensitivity to confinement of the outer leaflet probe in both nanodomain-rich and nanodomain-poor populations of Px units (Figure 3C,D). Similar differences in *D*_*av*_ for YFP-GL-GPI and PM-EGFP were observed by super-resolution SPT on RBL cells (63). YFP-GL-GPI also exhibits substantially higher values than PM-EGFP for *τ*_*0,av*_ and *Slope*_*av*_ (Figure 4E,F). The higher level of nanoscale confinement (*τ*_*0,av*_) may be due to more highly ordered nanodomains in the outer leaflet, consistent with slower *D* values and as suggested by comparing lipid compositions in both leaflets (85, 86). The higher apparent microscale viscosity (*Slope*_*av*_) may further reflect, for example, the thick glycocalyx on the outer leaflet that YFP-GL-GPI diffuses through in Ld-like regions (87). CytoD treatment has no effect on *τ*_*0,av*_, while the *Slope*_*av*_ increases markedly (Table 1, Figures 4C-E and S4b), suggesting that the largest changes of YFP-GL-GPI interactions to increase the apparent viscosity occur outside of Lo-like nanodomains.

Notably, PM-EGFP, Lyn-EGFP, and YFP-GL-GPI, exhibit *F*_*slow*_ values in the range of 0.63 – 0.72, independent of their *D*_*slow*_ and *D*_*av*_ values, which is consistent with previous measurements of ordered character in the plasma membrane of RBL cells by electron spin resonance, fluorescence anisotropy, and fluorescence imaging (88–90). In contrast, EGFP-GG exhibits *F*_*slow*_ = 0.41, consistent with this Ld-preferring probe being the least susceptible to confinement. Similar high coverage of ordered regions is also reported for other cell types (91, 92).

### Diffusion of TM proteins is influenced primarily by protein-based interactions

We tested AF488-IgE-FcεRI and other TM proteins with different numbers of TM segments and different preferences for Lo-like environments as evaluated by a variety of criteria (45, 76). Not surprisingly, the *D*_*av*_ of these protein probes are substantially slower than the lipid-anchored probes, and those with a single TM segment (YFP-GL-GT46, EGFR-EGFP) diffuse somewhat faster than those with four (AcGFP-Orai1) or seven (AF488-IgE-FcεRI) TM segments. All TM probes show *τ*_*0,av*_ values substantially higher than the lipid-anchored probes indicating confinement by protein complexes in or out of nanodomains (Table S2B). These confining interactions appear to be highly protein dependent as suggested by our results that probes with one TM segments show the smallest *τ*_*0,av*_, but the probe with seven TM segments exhibits a lower value than that with four TM segments. The *Slope*_*av*_ value, averaging over all interactions within the micrometer scale, is highest for the probe with seven TM segments and lowest for the probe with four TM segments, and these two values bracket the *Slope*_*av*_ values for all of the lipid-anchored probes in inner and outer leaflets. Overall, our results indicate that diffusion of TM protein probes is not dramatically influenced by Lo-like nanodomains in the resting plasma membrane but is most likely retarded predominantly by interactions with other proteins that depend on the biological and physical chemistry of the specific TM probe monitored.

### Concluding remarks

Statistical analyses of *D* distributions obtained from ImFCS has proven valuable for discerning weak interactions underlying plasma membrane heterogeneity in the poised, steady-state, as sensed by distinctive probes and as modulated by the actin cytoskeleton. This simple analytical framework can be readily adopted in other spatially resolved fluorescence fluctuation methods (48, 49, 93–97). In the context of mast cell activation, future studies will build on the foundation established here to examine stimulated signaling as mediated by the participating proteins within the environment of the responding membrane. A key step will be to quantitatively address the essential roles played by weak, lipid-based interactions in transmembrane signaling initiated by antigen-driven coupling of IgE-FcεRI with Lyn kinase anchored to the membrane inner leaflet (23).

## MATERIALS AND METHODS

### Reagents

Maximum essential medium (MEM), Opti-MEM, Trypsin-EDTA (0.01%) and gentamicin sulfate were purchased from Life Technologies (Carlsbad, CA). Fetal Bovine Serum (FBS) was purchased from Atlanta Biologicals (Atlanta, CA). Alexafluor 488 (AF488) NHS ester (Invitrogen) was used to label monoclonal anti-DNP immunoglobulin E (IgE) as described previously (77). Cytochalasin D (CytoD) and phorbol 12,13-dibutyrate (PDB) were obtained from Sigma-Aldrich (St. Louis, MO). Stock solutions of PDB and CytoD were prepared in DMSO and stored at −80°C.

### Cell culture, transfection, and labeling

RBL-2H3 cells (for brevity, RBL cells) were cultured in growth medium (80% MEM supplemented with 20% FBS and 10 mg/L gentamicin sulfate) at 37°C and 5% (v/v) CO_2_ environment. Cells in a confluent 75 cm^2^ flask were washed once with 2 mL Trypin-EDTA, trypsinized with 2 mL Trypin-EDTA for 5 min at 37°C and 5% (v/v) CO_2_ environment. Following trypsinization, cells were harvested in growth medium. About 20,000 cells were homogeneously spread in a 35 mm MatTek dish (Ashland, MA) containing 2 mL growth medium and allowed to grow overnight. MatTek dishes containing the adhered cells were transfected using FuGENE HD transfection kit (Promega). Typically, 0.5-1 μg of plasmid DNA and 3 μL FuGENE/μg DNA were used per dish. For transfection, plasmid DNA and FuGENE were first mixed in 100 μL Opti-MEM medium and incubated at room temperature for 15 min. Next, MatTek dishes containing cells were washed with 1 mL Opti-MEM and covered with another 1 mL Opti-MEM. The DNA/FuGENE complex was spread evenly over the cells and incubated for 1 hour, followed by incubation with pre-warmed PDB (1 mL, 0.1 μg/mL) for 3 hours at 37°C in 5% (v/v) CO_2_ environment. Finally, 2 mL of culture medium was added to each MatTek dish after discarding Opti-MEM. The transfected cells are allowed to grow for 18-22 hours at 37°C in 5% (v/v) CO_2_ environment before imaging. For imaging, the cells were washed twice with Buffered Salt Solution (BSS) solution (135 mM NaCl, 5.0 mM KCl, 1.8 mM CaCl_2_, 1.0 mM MgCl_2_, 5.6 mM glucose, and 20 mM HEPES; pH 7.4) and imaged in fresh BSS. The plasmids used in this study encode the following proteins: Lyn-EGFP (98), PM-EGFP (59), EGFP-GG (59), YFP-GL-GPI (65), AcGFP-Orai1 (99), EGFR-EGFP (100) and YFP-GL-GT46 (65).

For labeling FcεRI receptors, the cells were washed twice with BSS followed by addition of a mixture of AF488-labeled (0.5 μg/mL) and unlabeled (1.5 μg/mL) IgE for 40 min at room temperature. The cells were washed twice with BSS and directly imaged in fresh BSS.

For CytoD treatment, fresh working solution of CytoD (1 μM) was prepared by diluting the stock solution in BSS before the experiment. The DMSO content in the final solution was < 0.1%. The cells were pre-incubated with 1 mL of 1 μM CytoD for at least 15 min at room temperature before imaging.

### ImFCS data acquisition and analysis

Fluorescently labeled ventral plasma membranes of RBL cells were imaged with a home-built total internal reflection fluorescence microscope (TIRFM) (DMIRB, Leica Microsystems, Germany) equipped with an oil immersion objective (PlanApo, 100×, NA 1.47; Leica Microsystems, Germany), a 488 nm excitation laser (Coherent, Santa Clara, CA), and an electron multiplying charge coupled device (EMCCD) camera (black illuminated Andor iXON 897DU, pixel size 16 μm, Andor Technology, Belfast, UK). The excitation laser beam was introduced and focused on the back focal plane of the objective by a pair of tilting mirrors and a dichroic mirror (ZT405/488/561/640rpc, Chroma Technology). The same set of mirrors was used to adjust the TIRF angle of the excitation beam to illuminate the ventral membrane. The fluorescence signal from the sample was recorded by the EMCCD camera after it passes through the same objective and the dichroic mirror and reflected to the camera chip after being filtered by an emission filter (ZET488/561m, Chroma Technology). The laser power was 50 μW before objective.

For ImFCS, a stack of 80,000 images from a region of interest (ROI) on the plasma membrane was recorded at an acquisition speed of 3.5 ms/frame. The ROI sizes were between 40×40 to 50×50 pixels (pixel size in the object plane = 160 nm) depending on the cell size. The images were acquired in ‘frame transfer’ mode with 10 MHz read-out speed and an EM gain of 300 (scale 6-300) was used. Image acquisition was done using Andor Solis software. The acquisition conditions were optimized following the protocols reported previously (50, 101). All measurements were performed at room temperature.

This raw image stack was further processed by a FIJI plug-in for ImFCS (Imaging_FCS 1.491, downloaded on October 1, 2016 from the laboratory website of Prof. Thorsten Wohland, National University of Singapore, Singapore). Raw Autocorrelation Functions (ACFs) were first computed after 2×2 binning of an image stack (Px unit) and then each ACF was fitted with one-component, two-dimensional Brownian diffusion model (Equation 1). This yields a map of lateral diffusion coefficient (*D*) with pixel dimension of 320 nm.

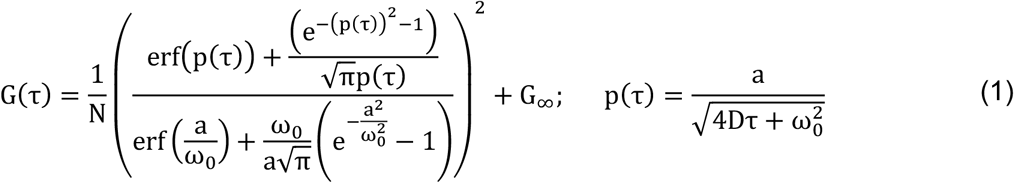

In the above equations, *G(τ)* is the ACF as a function of lag time (*τ*), *N* is the number of particles within the observation area, *D* is the lateral diffusion coefficient at the Px unit, *a* is the length of pixel in the object plane, *ω*_0_ is the point spread function (PSF) of the microscope, *A*_*eff*_ is the effective observation area, which is determined by the convolution of pixel area (*a^2^*) and point spread function, *G*∞ is the convergence value of *G(τ)* at very large lag time. We used *N, D* and *G*∞ as fit parameters, and *ω*_0_ was experimentally determined using the method described elsewhere (53).

The *τ*_*0*_ map was created by analysis spot variation FCS (svFCS) on small sub-regions (8×8 pixels) within the same raw stack (60). For this, diffusion times (*τ*_D_) were first determined for four different observation area (*A*_*eff*_) sizes generated by 2×2, 3×3, 4×4 and 5×5 binning within each 8×8 pixels sub-regions. The *A*_*eff*_ thus obtained were 0.42 μm^2^, 0.57 μm^2^, 0.78 μm^2^ and 1.05 μm^2^. Since fitting of ACF in ImFCS directly gives *D* value, the *τ*D values were determined by dividing *A*_*eff*_ with the corresponding *D* values. The plot of *τ*_*D*_ against *A*_*eff*_ for each sub-region was fitted with a straight line (Equation 2) to get the *τ*0 value as the intercept while the *Slope* of the fit is the inverse of effective long-range diffusion coefficient (1/*D*_*eff*_). There are 36 sub-regions (8×8 pixels) in a 50×50 pixels raw stack and thus the corresponding *τ*_*0*_ map will have 36 pixels. The pixel dimension of *τ*_*0*_ and *Slope* maps is 1.28 μm (8×160 nm).

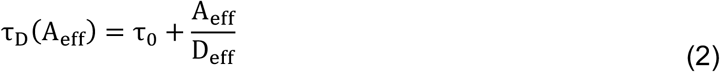

### Fitting of cumulative distribution functions (CDFs) of D

The *D* values obtained in Px units (Equation 1) from multiple cells for a given condition are grouped to create their respective distribution. First, cumulative frequencies for each *D* value were determined in ascending order, which was then plotted against corresponding *D* values to generated normalized CDF of *D* using Igor Pro (Version 7 and 8; WaveMetrics, OR, USA). This CDF was fitted with the following models (Equations 3 and 4 for one- and two-component models, respectively).

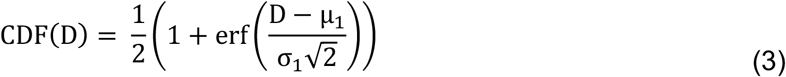

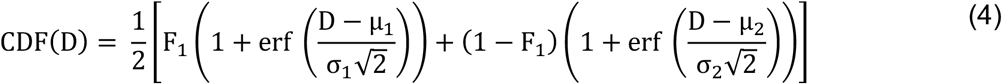

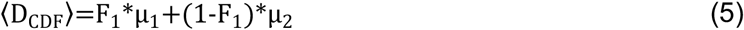

In the above equations, *μ*_1_ and σ_1_ are the mean and standard deviation of the first component while *μ*_2_ and σ_2_ are the mean and standard deviation of the second component, *F*_1_ is the fraction of first component and (1− *F*_1_) is the fraction of the second component of *D* distribution. The best fitting model (Equation 3 or 4) and goodness of fit were determined by reduced chi-squared values. We also compared the periodicity of the residual plots for both models. If fitting with one-component model yields residual plot that strongly oscillates around zero and this periodicity largely disappears in the residual plot for the fitting with two components, along with strong reduction of reduced chi-squared values, we accept a two-component model. A three-component model did not improve the quality of fitting in any case and therefore was not considered. The weighted average of *D*, <*D*_*CDF*_>, is given by Equation 5.

The component with smaller mean value, i.e., min {μ_1_, μ_2_} = *D*_*slow*_, is defined as representing the slower diffusing population, and the component with larger mean value, i.e., max {μ_1_, μ_2_} = *D*_*fast*_, is defined as representing the faster diffusing population. According to our working model for lipid-anchored probes (Appendix schemes A1-A3), *D*_*slow*_ represents probes diffusing in nanodomain-rich Px units, and *D*_*fast*_ represents probes diffusing in nanodomain-poor Px units.

## Abbreviations

ImFCS: Imaging Fluorescence Correlation Spectroscopy
svFCS: spot-variation Fluorescence Correlation Spectroscopy
CDF: Cumulative Distribution Function
GG: geranyl-geranyl lipid anchor
PM: minimal palmitoyl-myristoyl lipid anchor of Lyn
GPI: glycosylphosphatidylinositol anchor

## ACKNOWLEDGEMENTS

We are grateful to Dr. Alice Wagenknecht-Wiesner for the help in preparing the constructs. We thank Prof. Thorsten Wohland (National University of Singapore, Singapore) for making the ImFCS software freely available and helpful discussions. This work is supported by National Institute of Health Grant R01GM117552. The content is solely the responsibility of the authors and does not necessarily represent the of official views of the NIH.

