## Supplementary Information for "Imaging FCS Delineates Subtle Heterogeneity in Plasma Membranes of Resting Mast Cells"

#### Figure S1

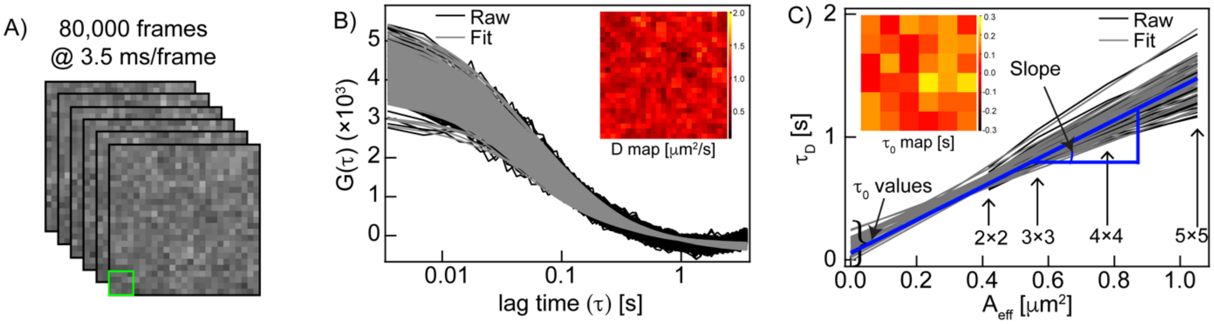


**Figure S1.** Principles of ImFCS data acquisition and analysis are illustrated with an EGFP-GG expressing RBL cell. **A**) Representative first few images of a 2×2 binned pixels (Px unit = 320×320 $nm^2$) image stack of 80,000 frames recorded with time resolution of 3.5 ms. The binning is done to obtain adequate signal-to-noise ratio for further data processing. **B**) Raw autocorrelation functions (ACFs) and respective fits for each Px unit using a single component Brownian diffusion model (Equation 1). Inset: spatial map of extracted diffusion coefficient (*D*) values at each Px unit of the image shown in (A). **C**) svFCS analysis is done on each Sv unit (8×8 pixels) to create svFCS plots (i.e., diffusion time ($\tau_D$) *vs* area ($A_{eff}$)). One representative Sv unit (1.28×1.28 $\mu m^2$) on the image stack is shown as green box in (A). $A_{eff}$ of various sizes are created by pixel binning within the Sv unit, as indicated on plot. svFCS analysis on all Sv units creates spatial map of $\tau_0$ (y-intercept) and *Slope* (e.g., blue line) values. Inset: spatial map of $\tau_0$ values determined from each Sv unit of the image shown in (A).

**Figure S2**

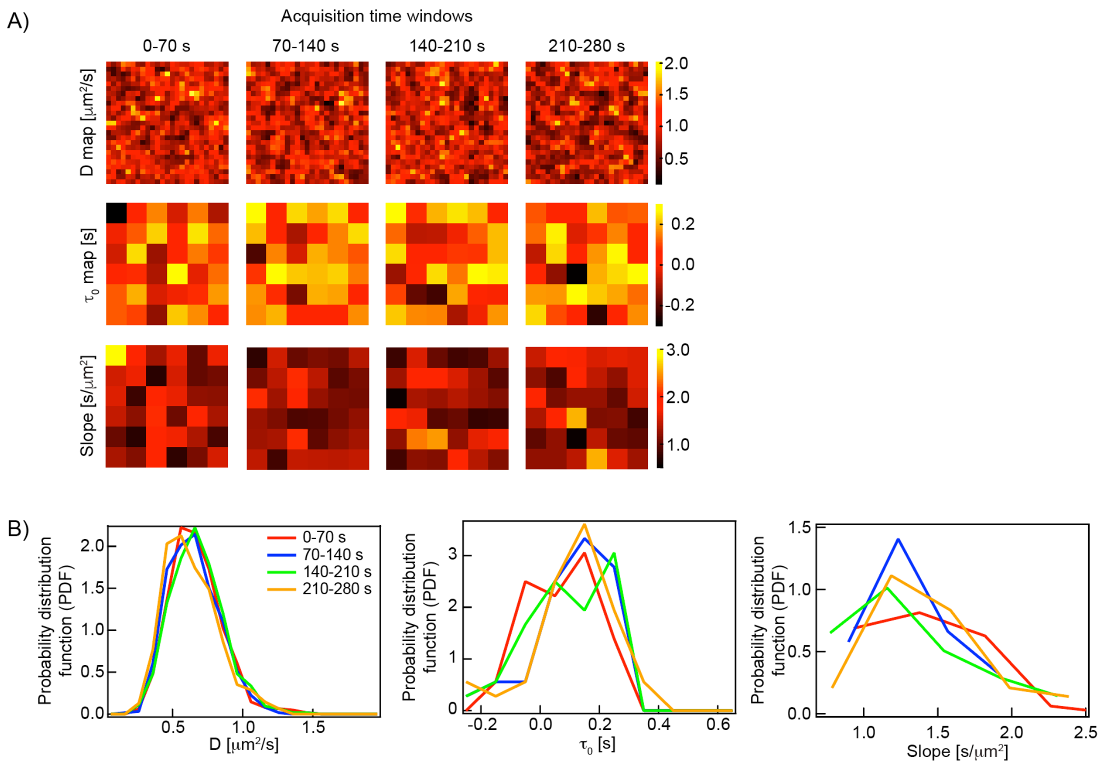


**Figure S2.** Temporal heterogeneity of membrane diffusion is demonstrated with a representative EGFP-GG expressing RBL cell. **A**) An 80,000 frames stack (280 s) is segmented into four equal 20,000 frames (70 s) time windows, and ACF and svFCS analyses are done (as in Figure S1) in each time window. Top, middle, and bottom rows provide the *D*, $\tau_0$ and *Slope* maps, respectively, from each time window. Pixel dimension of the *D* map is 320 nm (Px unit) and of $\tau_0$ and *Slope* maps is 1.28 μm (Sv unit). The locations of very high (yellow pixels) and very low (black pixels) values of these parameters fluctuate across the time windows indicating the dynamic nature of local diffusion. **B**) The *D*, $\tau_0$ and *Slope* values shown in the individual maps in (A) are translated into probability distribution functions, which show the shape and characteristics of these parameters remain roughly similar across all time windows.

#### *A large number of data points allow two components to be extracted from fitting of D CDFs*

Using the fitting strategy described in the Methods section (Equations 3 and 4), we observed that $\mu_1$ and $\mu_2$ values extracted from two-component fit are often very close with overlapping error bars (standard deviations) (Table 1). For instance, the CDF of *D* of EGFP-GG has two components: $D_{fast}$ = 0.66 ± 0.19 μm$^2$/s (59%) and $D_{slow}$ = 0.61 ± 0.09 μm$^2$/s (41%); the difference between the two components is 8% ($D_{fast}/D_{slow}$ = 1.08), which is the minimum among all probes reported here (summarized in Table 1). Therefore, we chose this distribution to test whether our pooled data are sufficiently robust statistically ($N_{Px}$ = $N_{total}$ = 10,527) to distinguish close populations. For this purpose, we generated a two-component Gaussian distribution (G) with variable total number of data points ($N_{total}$ = 10, 100, 500, 1000, 5,000, and 10,000) such that G

= 0.4*$G_1$ + 0.6*$G_2$. Here, $G_1$ and $G_2$ are two single-component Gaussian distributions. The mean ($\mu_1$) and standard deviation ($\sigma_1$) of $G_1$ were 0.66 and 0.19 while those of the $G_2$ were 0.61 ($\mu_2$) and 0.09 ($\sigma_2$). The generation of random numbers, construction of CDFs and fitting were performed in Igor Pro (Version 7 and 8).

Figure S3 shows that at least 500 data points are needed to construct a smooth distribution. However, fitting does not yield the correct values of the individual components for $N_{total}$ = 500 and the reduced chi-squared values do not improve significantly from one-component to two-component fit (Table S1). This is also observed in noisy residual plots (green in Figure S3B, C). At least 5,000 data points are required to reliably estimate the two populations (Table S1 and Figure S3D). Our experiments generate more than 5,000 experimental *D* values for all membrane probes tested (Table 1, main text), such that our analyses do not over-fit the experimental CDFs of *D.*

Generally, we choose two-component fit model if all of the following criteria are satisfied:

1. Residual plot of one-component fit is strongly periodic while no obvious periodicity is observed in the residual plot of two-component fit.

2. Reduced chi-squared of two-components fit is more than 10 times smaller than that of one-component fit.

3. Fit parameters after two-component fit are such that $D_{fast}/D_{slow} \geq 1.1$.

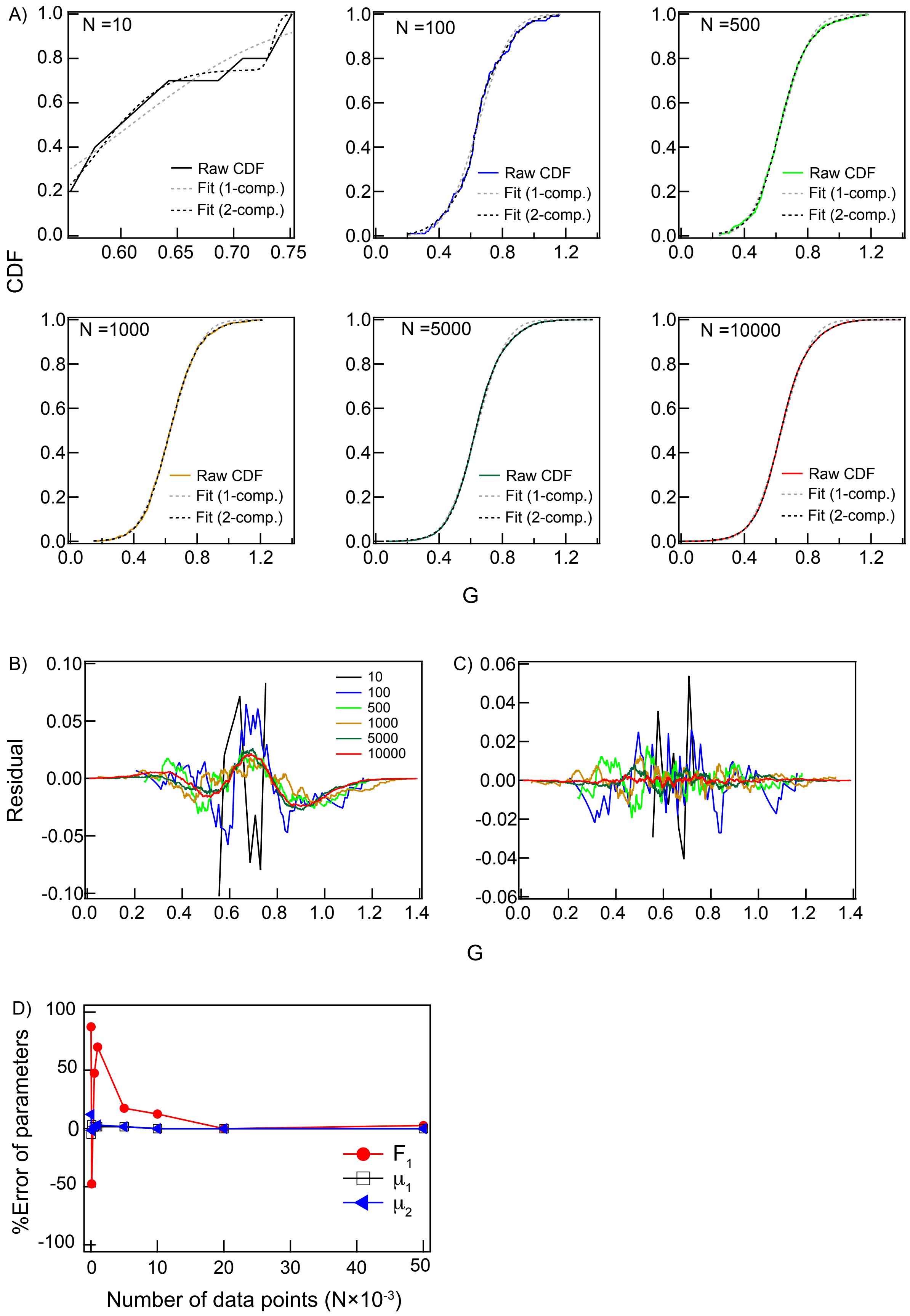

A)
N =10
N =100
N =500
N =1000
N =5000
N =10000
Raw CDF
Fit (1-comp.)
Fit (2-comp.)
CDF
G
B)
C)
Residual
10
100
500
1000
5000
10000
D)
%Error of parameters
Number of data points (N×10^-3)
F1
μ1
μ2

**Figure S3.** Simulated CDFs with two-component diffusion coefficients require thousands of data points to be accurately fit. **A**) Gaussian distributions with different number of points are constructed by combining two one-component Gaussian distributions with characteristic mean and standard deviations (First distribution ($G_1$): mean ($\mu_1$) = 0.61, standard deviation ($\sigma_1$) = 0.09, weight ($F_1$) = 40%; second distribution ($G_2$): mean ($\mu_1$) = 0.66, standard deviation ($\sigma_1$) = 0.19, weight ($F_1$) = 60%). Total number of points that are used to construct the final distribution is given in the individual panels. The colored solid lines are raw data; grey and black dotted lines are one-component and two-component fits, respectively. **B**) Residual plot of all distributions shown in (A) for one-component fit clearly showing oscillation. **C**) Residual plot of all distributions shown in (B) for two-component fit. **D**) Error of the estimated values of $F_1$, $\mu_1$ and $\mu_1$ as a function of number of input data points.

### Table S1

Fitting results of simulated two-component Gaussian distributions presented in Figure S3

| **Number of data points** | **One-component fit (Equation S3)** | | | **Two-component fit (Equation S4)** | | | | | | |
|---|---|---|---|---|---|---|---|---|---|---|
| | $\mu_1$ | $\sigma_1$ | $\chi^2_{red}$ | $\mu_1$ | $\sigma_1$ | $\mu_2$ | $\sigma_2$ | $F_1$ | $F_2$ | $\chi^2_{red}$ |
| 10 | 0.61 | 0.10 | 0.040 | 0.74 | 0.06 | 0.58 | 0.05 | 0.25 | 0.75 | 0.008 |
| 100 | 0.65 | 0.16 | 0.061 | 0.65 | 0.20 | 0.63 | 0.03 | 0.79 | 0.21 | 0.012 |
| 500 | 0.63 | 0.15 | 0.103 | 0.67 | 0.22 | 0.62 | 0.11 | 0.41 | 0.59 | 0.020 |
| 1000 | 0.63 | 0.15 | 0.081 | 0.68 | 0.21 | 0.62 | 0.12 | 0.32 | 0.68 | 0.015 |
| 5000 | 0.64 | 0.15 | 0.648 | 0.67 | 0.19 | 0.62 | 0.10 | 0.53 | 0.47 | 0.028 |
| 10000 | 0.64 | 0.15 | 1.152 | 0.66 | 0.19 | 0.61 | 0.09 | 0.55 | 0.45 | 0.007 |

**Figure S4**

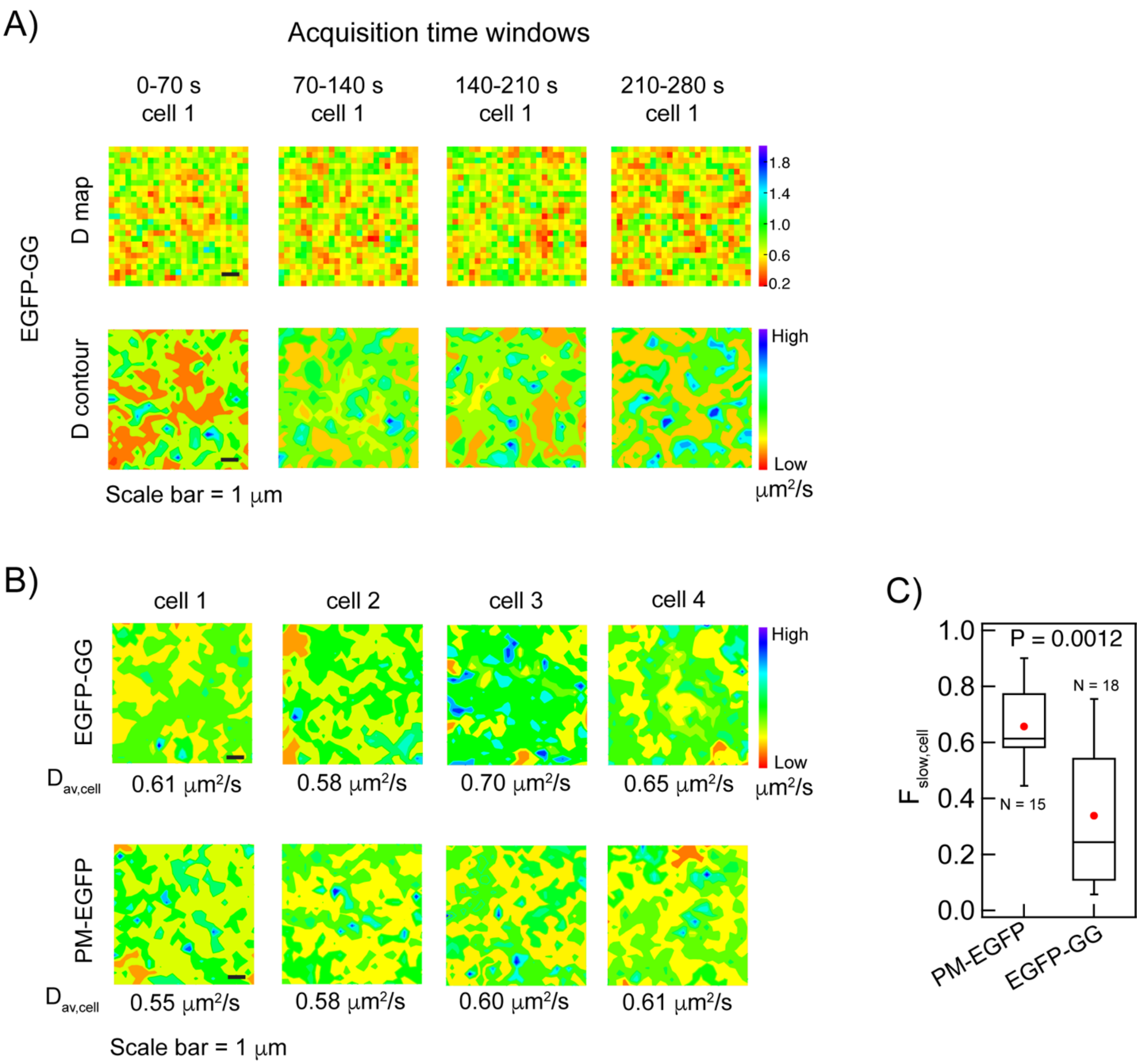


**Figure S4.** Contours of *D* value distributions for single cells show dynamic heterogeneity and distinguish EGFP-GG from PM-EGFP. **A**) Contours of the *D* maps presented in Figure S2A (70 s acquisition time windows) showing the temporal heterogeneity of the diffusion distribution across the plasma membrane of one representative cell. The color scheme differs from Figures S1 and S2 for better visualization of the contours. **B**) Steady-state contours of *D* maps (full 280 s acquisition) for four representative cells expressing either EGFP-GG or PM-EGFP. The contours show relative differences in the diffusion of these two probes. In addition, they show spatial heterogeneity of diffusion among different cells even though the arithmetic average of *D* values obtained from the individual cells ($D_{av,cell}$) are similar. The contours also demonstrate relatively faster diffusion as major component for EGFP-GG (higher coverage of green to cyan areas) and relatively slower diffusion as major component for PM-EGFP and EGFP-GG (higher coverage of red to yellow areas) for individual cells in agreement with the CDF analysis of the pooled *D* values from multiple cells (Table 1, main text). **C**) Box plots show the value of $F_{slow,cell}$ calculated from contour maps of N cells based on the fraction of total image area covered by slower diffusion (red to yellow areas). This statistical analysis was performed using Mann-Whitney test, and resulting P values are provided in individual panels; the box height corresponds to 25th to 75th percentile and the error bars represent 10th to 90th percentile of the entire data set. The mean and median values are represented as solid circle and bar, respectively, located inside the box. These mean values for single cells expressing EGFP-GG or PM-EGFP agree well with CDF analysis from multiple cells (Table 1).

### Figure S5

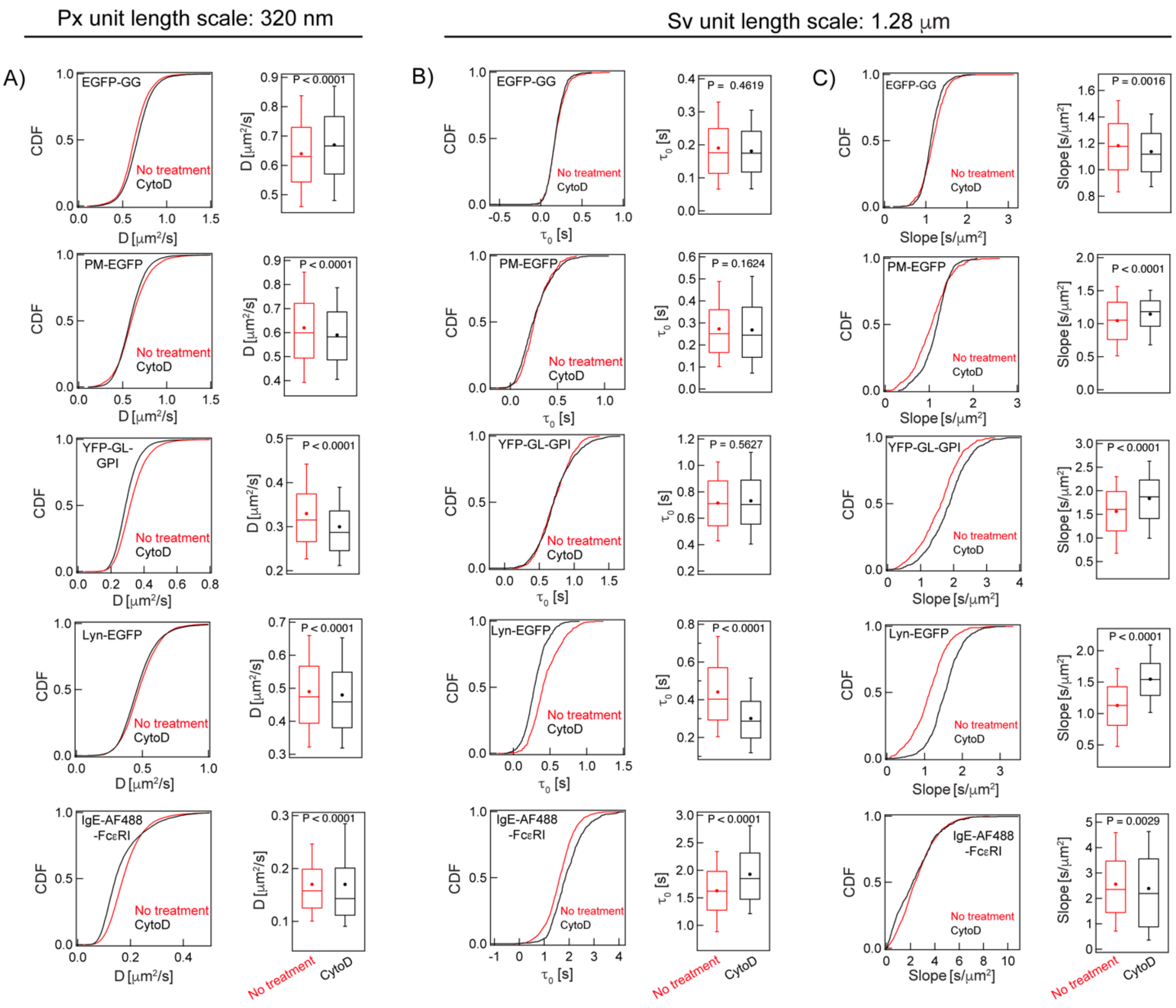


**Figure S5.** CDFs of diffusion properties for inner-leaflet, outer-leaflet and TM probes change variably when actin polymerization is inhibited. Probes evaluated are depicted in Figure 1B (main text). **A**) *D* distributions, **B**) $\tau_0$ distributions and C) *Slope* distributions without (black) and with (red) CytoD treatment. Box plots of values in CDFs are shown to the right; the box height corresponds to 25th to 75th percentile and the error bars represent 10th to 90th percentile of the entire data set. The mean and median values are represented as solid circle and bar, respectively, located inside the box; this statistical analysis was performed using Mann-Whitney test, and resulting P values are provided in individual panels. The mean values of CDFs are provided as $D_{av}$, $\tau_{0,av}$ and $Slope_{av}$ in Table 1 (main text). The number of data points used to construct the CDFs and corresponding Box plots are provided as $N_{Px}$ (for *D*) and $N_{sv}$ (for $\tau_0$ and *slope*) in the Table S2 below (also in Table 1).

**Table S2**

Numbers of data points used to create the plots in Figure S5

| **Probes** | **Membrane association** | **Treatment** | **$N_{Px}$/$N_{sv}$ (No. of cells)** |
|---|---|---|---|
| EGFP-GG | Inner leaflet, Ld-preferring | No | 10527/648 (18) |
| | | CytoD | 9366/539 (16) |
| PM-EGFP | Inner leaflet, Lo-preferring | No | 9375/540 (15) |
| | | CytoD | 12432/720 (20) |
| YFP-GL-GPI | Outer leaflet, Lo-preferring | No | 12892/719 (21) |
| | | CytoD | 11074/661 (19) |
| Lyn-EGFP | Inner leaflet, Lo-preferring | No | 10000/575 (16) |
| | | CytoD | 12904/753 (21) |
| AF488-IgE-FcεRI | TM, 7 TMD | No | 27707/1644 (46) |
| | | CytoD | 10624/558 (17) |

**Figure S6**

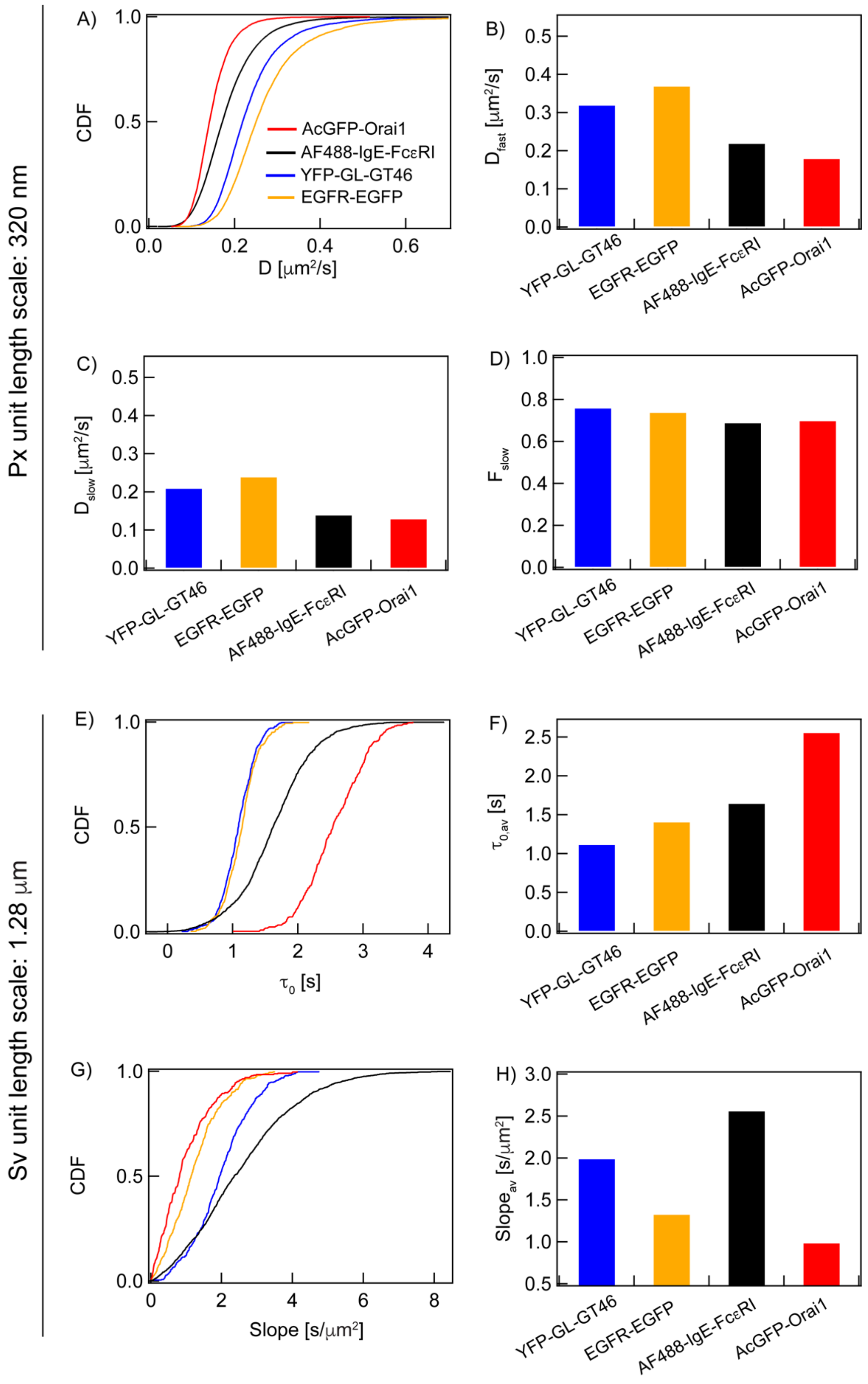
Px unit length scale: 320 nm
A)
CDF
D [μm²/s]
AcGFP-Orai1
AF488-IgE-FcεRI
YFP-GL-GT46
EGFR-EGFP
B)
D_fast [μm²/s]
C)
D_slow [μm²/s]
D)
F_slow
YFP-GL-GT46
EGFR-EGFP
AF488-IgE-FcεRI
AcGFP-Orai1
Sv unit length scale: 1.28 μm
E)
CDF
τ_0 [s]
F)
τ_0,av [s]
G)
CDF
Slope [s/μm²]
H)
Slope_av [s/μm²]

**Figure S6.** CDFs and extracted *D*, $\tau_{0,av}$, and $Slope_{av}$ values differ for transmembrane (TM) probes expressed in resting RBL cells: AcGFP-Orai1 (red), AF488-IgE-FcεRI (black), EGFP-EGFP (orange) and YFP-GL-GT46 (blue). **A**) *D* CDFs of the probes. **B-D**) Fits of *D* CDFs with Equation 4 to yield $D_{fast}$, $D_{slow}$ and $F_{slow}$. **E-F**) $\tau_0$ CDFs and $\tau_{0,av}$ values of TM probes. G-H) *Slope* CDFs and $Slope_{,av}$ values of TM probes. Numerical values of all extracted parameters with defined errors are provided in Table 1 (main text).

### APPENDIX. PLASMA MEMBRANE 'SENSED' BY DIFFUSING PROBES DEPENDS ON STRUCTURAL INTERACTIONS

In this section, we illustrate the distributions of diffusion coefficients, *D,* expected for lipid-anchored probes diffusing through the heterogeneous plasma membrane. As in our experiments, the plasma membrane ROI is divided into roughly 64 Px units of 320×320 $nm^2$, which is the resolution of our current ImFCS measurements. A *D* value is determined for each Px unit, and these are compiled into a probability distribution function (PDF) or a cumulative distribution function (CDF). The *D* of a probe moving through a Px unit will depend on its structural interactions therein. As described in the Introduction, we take the basic view that the plasma membrane comprises Lo-like proteolipid nanodomains connected by Ld-like regions. Lipid-anchored probes are dynamically confined within the nanodomains depending on their level of interactions; the diffusion of acyl chains that are more Lo-like (e.g., PM-EGFP) will be retarded more than those that are more Ld-like (e.g., EGFP-GG). We assume that all probes diffuse freely in the Ld-like regions, and any interactions there are less than those within the nanodomains and will manifest in an apparent, micrometer-scale viscosity. We include the possibility of additional protein-based interactions inside and outside these nanodomains for probes with functional protein modules (e.g., Lyn-EGFP). In general, a probe partitioning into the nanodomains will be transiently retained and thus will diffuse more slowly inside compared to outside nanodomains.

#### <u>One- and two-component D PDFs/CDFs are observed in ImFCS depending on relative coverage by nanodomains in Px units</u>

In schemes A1a-c ordered nanodomains are differentially distributed (left side): a) all Px units are filled with nanodomains (no Ld-like regions), b) all Px units are devoid nanodomains (all Ld-like), and c) 70% of the Px units (i.e., 45 out of 64 Px units) are filled with nanodomains. Although the dimensions of individual nanodomains in the plasma membrane are on the order of 10 nm, much less than 320 nm, for purposes of illustration we take the area of a nanodomain as equal to the area of a Px unit. The blue trajectories indicate the diffusion trajectory of a probe (e.g., PM-EGFP) as it diffuses through the nanodomains, if any, and across the Px units.

The observed D values for a probe diffusing in these three scenarios will be different. Scheme A1a will yield a single slower value and Scheme A1b will yield a single faster value, as represented by the respective PDFs for these two schemes (right side).

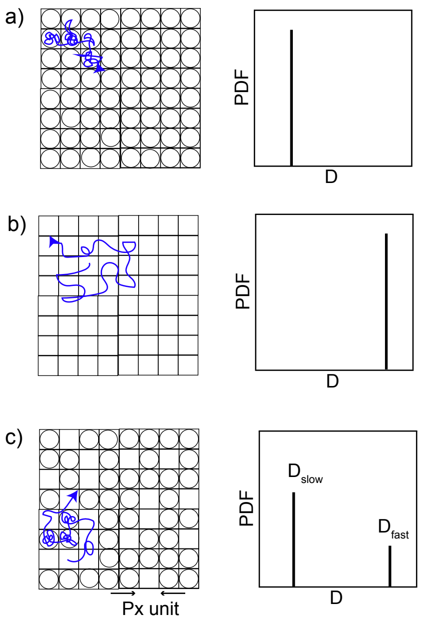


**Scheme A1**

In scheme A1c, 45 of 64 Px units (~70%) are filled with nanodomains while the other 19 Px units are entirely Ld-like. Correspondingly, 70% of the $D$ values will be singly slow (the same value as in Scheme A1a), and 30% will be singly fast (the same value as in Scheme A1b). The observed PDF of $D$ values from the entire ROI will thus have two components with values of $D_{\text{slow}}$ (associated with nanodomain-rich Px units) and $D_{fast}$ (associated with nanodomain-poor Px units). The fraction of Px units with $D_{slow}$, ($F_{slow}$) will be 0.70.

Schemes A1a-c simply demonstrate the capacity of ImFCS to reveal the presence of nanodomains within Px units from $D$ PDFs measured for a probe such as PM-EGFP. However, in a complex plasma membrane the relative area covered by the nanodomains may vary significantly, as illustrated in Scheme A2, which does not yield distinct, singular values of $D$ for a diffusing probe. Rather, PDFs represent overlapping populations of relatively nanodomain-rich and nanodomain-poor Px units. Although these populations as experienced by the diffusing probe may be separately delineated, they blur into Gaussian shapes that are distributed around the most probable values. As demonstrated in Figures 2 (main text) and S3, the robust experimental statistics of the current ImFCS study (~10,000 $D$ values from Px units in multiple cells) allow precise fitting with one- or two-component Gaussian models. These measurements can distinguish a maximum of two underlying populations from experimental $D$ PDFs, if $D_{fast}/D_{slow} \geq 1.1$, as illustrated in the right panel of Scheme A2. Thus, a two-component PDF (dotted line) can be resolved by ImFCS if $D_{fast}$ is at least 10% greater than $D_{slow}$. Otherwise, we reliably extract a single component (solid line) with a shape and magnitude representing the weighted average of any underlying components. Similarly, for a two-component $D$ PDF, $D_{slow}$ represents averaged $D$ values of potentially multiple populations of nanodomain-rich Px units, and $D_{fast}$ represents all sub-populations of nanodomain-poor Px units.

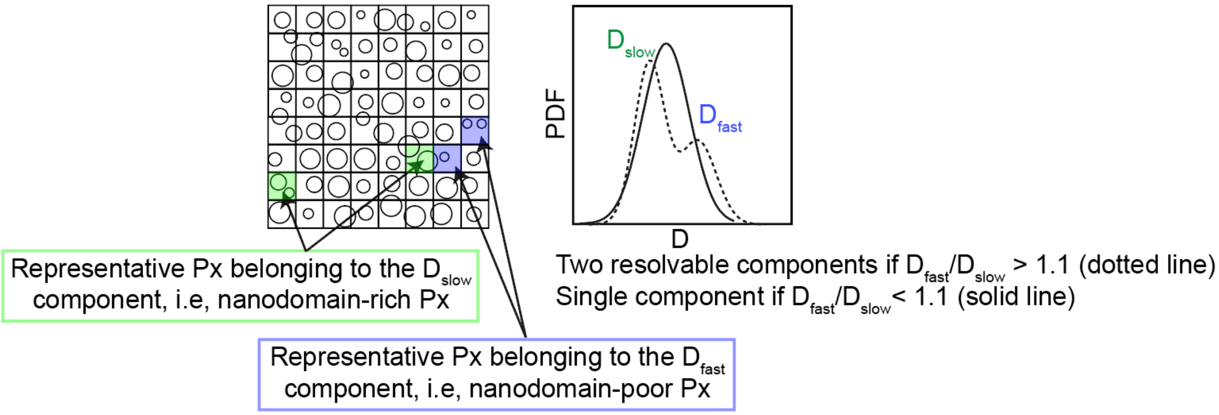


**Scheme A2**

**Integrated view of plasma membrane heterogeneity as sensed by structurally distinct membrane probes.** The same steady-state organization of the nanodomains in the plasma membrane will be reflected differentially in *D* PDFs determined experimentally for different probes, depending on the strength of interactions between the probes and nanodomains. The diffusion of a particular probe may be delayed to the same extent by two chemically distinct nanodomains, for example a different mix of proteolipid constituents. Also, two different types of membrane probes may interact with same nanodomain differently. For example, we expect lipid-anchored probes with Lo-preferring acyl chains (e.g., PM-EGFP) to yield a *D* PDF different from those with Ld-preferring acyl chains (e.g., EGFP-GG). In addition to lipid-based partitioning we consider protein-based interactions a given probe might undergo. For example, Lyn-EGFP has same lipid anchor as PM-EGFP but also possesses cytosolic protein modules which are likely to participate in protein-protein interactions both inside and outside of the proteolipid nanodomains. With ImFCS we can distinguish a maximum of two components, but we expect *D* PDFs for each probe diffusing within the same membrane milieu to be distinctive and reflect its particular set of interactions. Similarly, we expect the effect of a given membrane perturbation (e.g., inhibition of actin polymerization) to be variable for different probes.

*Lipid-anchored probes*: Trends expected for the three inner-leaflet probes can be integrated into a unified model (Scheme A3, left panel). PM-EGFP, which shares the Lo-preferring, saturated lipid anchors of Lyn-EGFP but not the protein modules, is confined less and diffuses relatively faster through the nanodomains due to lack of protein-based interactions. EGFP-GG possesses Ld-preferring, unsaturated lipid anchors and no interactive protein modules. Relative *D* PDFs of PM-EGFP and Lyn-EGFP and EGFP-GG are shown in Scheme A3, right panels. The experimentally observed two-component *D* PDFs for all three of these probes as well as the trend in absolute values of $D_{av}$ (Lyn-EGFP < PM-EGFP < EGFP-GG) (see Table 1, main text) are consistent with this view. The same reasoning holds for the diffusion of lipid-anchored probes in the outer leaflet. However, the physical characteristics of the nanodomains in the outer leaflet are likely to differ from those in the inner leaflet due to asymmetric lipid composition and differences in the membrane-proximal interactions. This is evident when the diffusion of YFP-GL-GPI and PM-EGFP, the Lo-preferring probes for outer and inner leaflet respectively, are compared (Table 1).

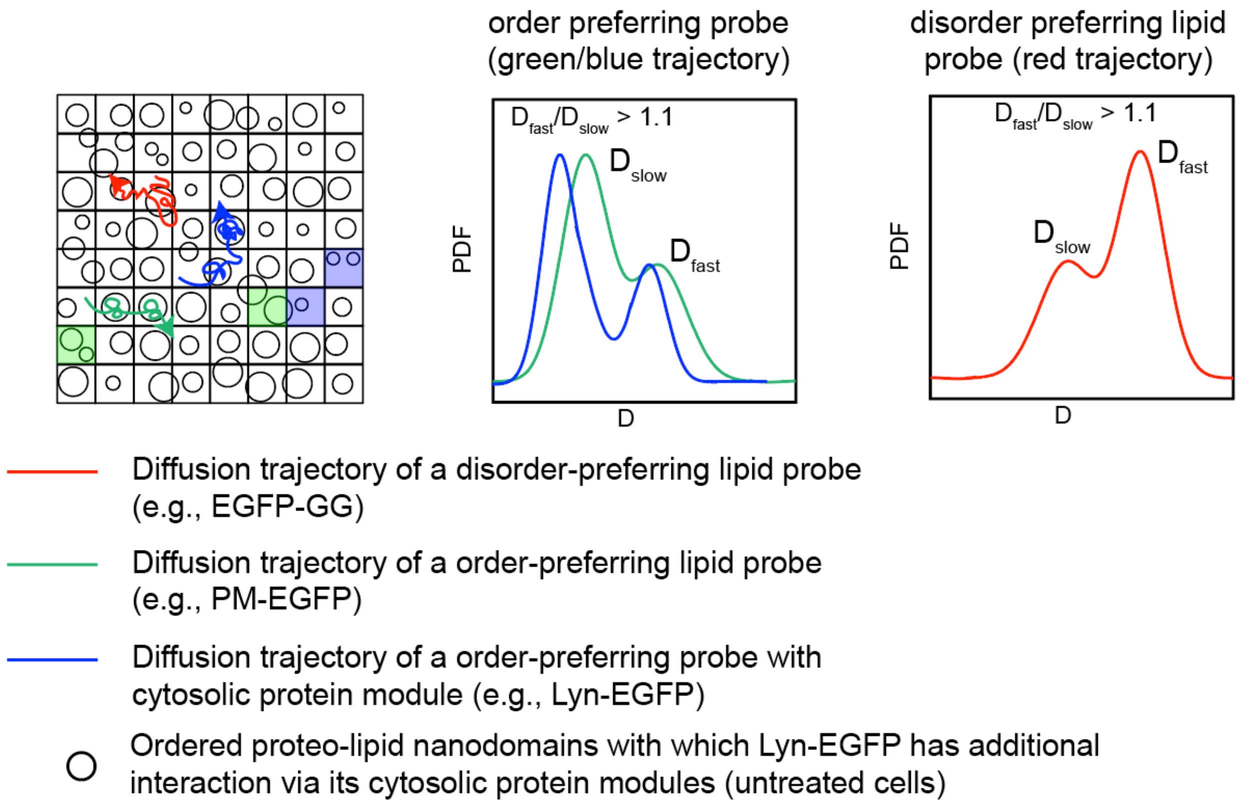


**Scheme A3:** Organization of plasma membrane inner leaflet as sensed by EGFP-GG, PM-EGFP and Lyn-EGFP.

*Transmembrane Probes:* The two-component distribution of *D* values observed for transmembrane probes (Table 1) can be explained by organization of membrane heterogeneity in the plasma membrane following the same principles described for lipid-anchored probes in Schemes A1-A3. However, the variety of interactions that TM probes are involved in are more complex than lipid-anchored probes and are likely to be predominantly protein-based rather than lipid-based. These would include protein-specific interactions via cytosolic and exoplasmic modules as well as interactions within the bilayer plane between the α-helix of the protein and membrane lipids. Because observed diffusion properties of the TM probes are not simply explained by their interactions with Lo-like proteolipid nanodomains, we define the Px units corresponding to $D_{slow}$ and $D_{fast}$ as interaction-rich and interaction-poor, respectively.
